# Measuring proteomes with long strings: A new, unconstrained paradigm in mass spectrum interpretation

**DOI:** 10.1101/282624

**Authors:** Arun Devabhaktuni, Niclas Olsson, Carlos Gonzales, Keith Rawson, Kavya Swaminathan, Joshua E. Elias

**Author notes:** Corresponding Author: Dr. Joshua Elias, Clark Center W300C, 318 Campus Drive, Stanford, CA 94305.

## Abstract

Thousands of protein post-translational modifications (PTMs) dynamically impact nearly all cellular functions. Mass spectrometry is well suited to PTM identification, but proteome-scale analyses are biased towards PTMs with existing enrichment methods. To measure the full landscape of PTM regulation, software must overcome two fundamental challenges: intractably large search spaces and difficulty distinguishing correct from incorrect identifications. Here, we describe TagGraph, software that overcomes both challenges with a string-based search method orders of magnitude faster than current approaches, and probabilistic validation model optimized for PTM assignments. When applied to a human proteome map, TagGraph tripled confident identifications while revealing thousands of modification types on nearly one million sites spanning the proteome. We expand known sites by orders of magnitude for highly abundant yet understudied PTMs such as proline hydroxylation, and derive tissue-specific insight into these PTMs’ roles. TagGraph expands our ability to survey the full landscape of PTM function and regulation.

## Introduction

Post translational modifications (PTMs) dynamically modulate the activity, conformation states, localization, interactions, abundance, and degradation of almost all proteins encoded by the human genome^1–4^, yet most remain poorly understood. PTM dysregulation has been linked to heart^5^, neurodegenerative^6^, and autoimmune^7^ diseases, cancer^8^, and countless other major health challenges^9^. Thus, characterizing PTMs’ identities, abundances, and regulation is an essential dimension for understanding overall protein function and disease etiology. However, mapping the full breadth of PTM identities and locations across the entire human proteome has remained intractable^10^.

Mass spectrometry is arguably the most robust technology capable of direct, unambiguous, and large-scale PTM measurement. It has provided transformative insight into the roles phosphorylation^11^, acetylation^12^, ubiquitylation^13^ – both singly and in combination^14^ -- have on cell biology. However, most global PTM studies have focused on modifications with optimized enrichment workflows^15^. Consequently, our current view of PTMs’ collective impact on the human proteome is heavily skewed towards a small fraction of the possible PTM landscape^16^.

Even without experimental enrichment, PTM-containing peptides are readily detected by routine tandem mass spectrometry (MS/MS) experiments^17, 18^, and are believed to comprise much of the “dark matter” in proteome datasets that consistently evades confident identification^19^.

Conventional sequence database search tools cannot identify modified peptides unless they are first anticipated by the researcher^20–22^. Search parameters including the number, kind, and frequency of PTMs are usually chosen to strike a difficult compromise: considering larger numbers of PTMs and other sequence variants is necessary for their identification, but doing so exponentially increases the time needed to interpret MS/MS datasets, and decreases the ability to distinguish correct from incorrect assignments^23^. To partially address this compromise, strategies have been proposed to constrain the number of proteins being searched, protease specificity rules, or the allowable types and numbers of PTMs^17, 18, 24–26^. In practice, these approaches only marginally decrease search times without clearly distinguishing correct from incorrect PTM assignments^27^. Therefore, most have not been demonstrated on large, proteome-scale datasets^23^

Here, we describe *TagGraph*, a powerful computational tool that addresses two principle challenges of searching very large sequence spaces. First, *TagGraph* leverages accurate *de novo* mass spectrum interpretations^28, 29^ to rapidly search millions of possible sequences for a match with an FM-index^30^ data structure. This highly efficient search method makes modern next-generation genome sequencing possible^31^, but has not been adapted to proteomics. By combining it with a graph-based string reconciliation algorithm, TagGraph rapidly searches MS/MS datasets without restrictions on number of proteins, PTMs, or protease specificity. This strategy achieves speeds orders of magnitude faster than prior algorithms because it considers exponentially more sequence possibilities without having to explicitly test each one against input spectra. Second, by replacing conventional “target-decoy” error estimation^32^ with a PTM-optimized probabilistic model, TagGraph accurately discovers and discriminates high-confidence peptide identifications even from such large search spaces. We demonstrate 40-fold more accurate false discovery rate estimation relative to target-decoy-dependent software.

Combined, these advances make large-scale, untargeted PTM proteomics possible. We demonstrate this new search capability on a recently published human proteome draft^33^. Our analysis reveals 2,576 modification types and 936,886 total modification sites spanning 13,791 proteins across 30 adult and fetal tissues (44,232 likely PTM sites across 5,576 proteins), and with accurately estimated false discovery rates commensurate with current proteomics standards. Simultaneously assaying such a large number of modifications substantially expanded the number of known PTM sites by orders of magnitude, particularly for those lacking biochemical enrichment techniques. Our analysis reveals quantitative, functionally relevant, differences in PTM stoichiometry between human tissues. We focus particularly on hydroxylation, demonstrating this un-enrichable PTM’s prevalence in the human proteome, its potential role in cancer, and its association with candidate enzymes. Through this analysis, we establish TagGraph as a paradigm shift for rapid proteome characterization, which promotes simultaneous, unbiased PTM identification.

## Results

### TagGraph: A new paradigm in fast, unrestricted proteome analysis

We developed TagGraph to address the compromise between search accuracy, depth, and speed proteomics researchers commonly face when searching large tandem mass spectra (MS/MS) datasets. With it, we can now accurately assign peptides bearing multiple unspecified post-translational modifications (PTMs) or amino acid substitutions. Conventional database search algorithms perform exponentially more comparisons between MS/MS spectra and peptide candidates as they consider new modification types. In contrast, TagGraph rapidly selects a very small number of candidate peptides from a sequence database through an efficient string matching and reconciliation procedure (Fig. 1a, Supplementary Fig. 1). In so doing, TagGraph effectively surveys very large sequence spaces that would be impractical to query using traditional database search engines.

**Figure 1.**
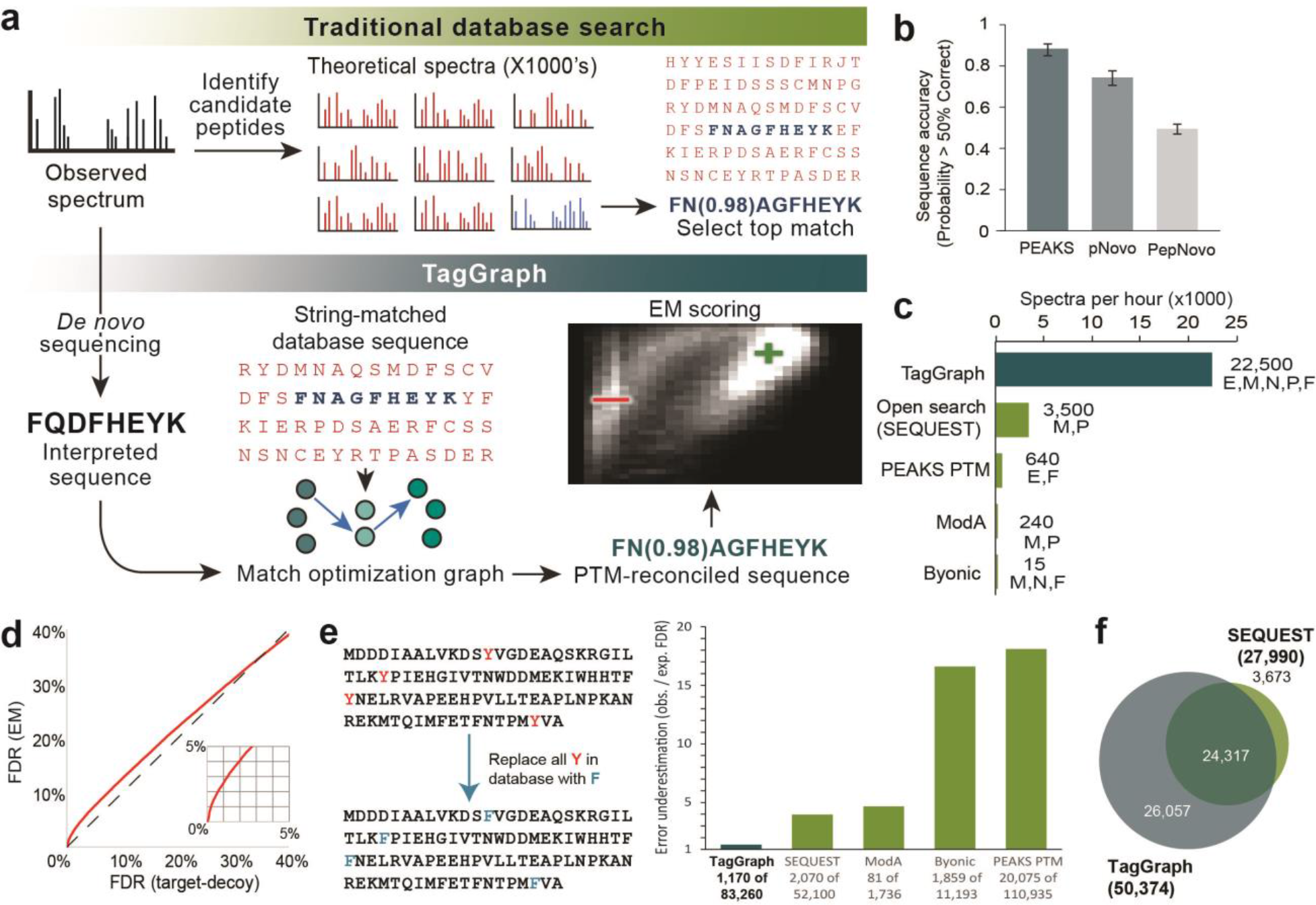
Through flexible string matching, TagGraph efficiently enables in-depth proteome characterization while controlling identification error. **a)** TagGraph workflow. Traditional database search engines compare an observed MS/MS spectrum to hundreds or thousands of peptide candidates. In contrast, TagGraph first extracts a candidate peptide sequence from high resolution MS/MS spectra by *de novo* sequencing to facilitate rapid indexed protein database searching, and sequence reconciliation. This process lets TagGraph consider an unlimited number of PTMs and amino acid substitutions. A small number of top-scoring sequence candidates are ultimately scored against the input spectrum using an EM-optimized probabilistic Bayesian network. **b)** The majority of *de novo*-interpreted high-resolution MS/MS spectra are mostly correct. The proportion of analyzed spectra interpreted with over 50% sequencing accuracy by PEAKS ^46^, pNovo ^84^, and PepNovo ^85^ on a data set of 168,391 MS/MS spectra derived from the A375 melanoma cell line. Error bars correspond to the standard deviation in accuracy over different fractions from this data set. ROC curves corresponding with these graphs are described in Supplementary Fig. 2b. **c)** TagGraph search times on the A375 data set were at least an order of magnitude faster than previously described unconstrained modification and iterative search strategies, even when the latter were given comparatively reduced search spaces (Table 1). Letters indicate which search space expansions were compatible with the search algorithm: E, no enzyme specificity; M, any possible modification; N, any number of modifications per peptide; P, all proteins in sequence database. F indicates that the algorithm estimates a false discovery rate from its identifications. **d)** Expectation Maximization-based false discovery rate estimation is generally consistent with, but more conservative than traditional target-decoy-based estimates when both are applied to TagGraph results. This is expected considering target-decoy’s inability to distinguish correct and incorrect modification annotations **e)** The human proteome sequence database was modified, substituting every tyrosine residue with a phenylalanine. The accuracy of PTM-specific false discovery rates was estimated based on substituted phenylalanine-containing peptides reported by each algorithm (Methods). Only TagGraph reported results with an empirically calculated FDR close to 1%. Open search^18^ was not included in this error rate comparison because it does not directly localize modifications to specific residue positions. **f)** By allowing unrestricted modifications to candidate peptides, TagGraph identified nearly twice as many unique peptide forms as SEQUEST, configured with common search parameters. Analogous to the proteoform concept^86^, we define unique peptides by the combination of the peptide’s amino acid sequence and any modifications made to them.

TagGraph leverages the speed of indexed string matching algorithms^34^ by first transforming complex, numeric mass spectra into discrete, unambiguous query strings using *de novo* peptide sequencing. *De novo* sequencing algorithms produce long, reasonably accurate sequence predictions from high resolution MS/MS spectra^29, 35^: we found these predictions were over 50% correct for nearly all interpretable MS/MS spectra (Fig. 1b, Supplementary Fig. 2)^29^.

Consequently, we reasoned that many *de novo* peptides should contain a sub-string that perfectly matches the true protein source of the observed MS/MS spectrum. The FM-index data structure^36^ was developed to facilitate this kind of search. TagGraph uses it to rapidly assemble a small number of candidate peptide matches from an arbitrarily large, pre-indexed sequence database with no restrictions on protease specificity, post-translational modifications, or sequence variants. These candidates are then reconciled against the input *de novo* sequence using a graph-based alignment algorithm that can discover and localize multiple PTMs and other sequence alterations that co-occur on a single peptide sequence without anticipating them *a priori* (**Supplementary Note 1**). Modification masses localized to individual amino acid positions within a peptide are cross-referenced with the Unimod resource^10^ to suggest the modification’s most likely identity based on mass and amino acid specificity. In this way, TagGraph effectively searches all possible sequence alterations on time scales commensurate with conventional database search tools.

Our strategy contrasts with prior approaches that extract many (>100) short sequence fragments (“tags”) from each input MS/MS spectrum to restrict protein candidates^37–39^. Although they consider fewer peptides per mass spectrum than conventional database search algorithms, they are subject to similar speed limitations if they consider large numbers of amino acid modifications and variants. Similarly, recent iterative approaches towards refining candidate proteins and modifications^40^, or non-specific database searches using very wide mass tolerances^18, 37^ are subject to these limitations: their ability to identify modifications requires comparison between spectra and large numbers of modified peptide candidates. As a result, these approaches are prone to infeasibly long search times. We measured the advantage of shifting this computational burden to TagGraph’s string matching and reconciliation procedure by comparing TagGraph’s execution time to four algorithms designed to consider greatly expanded search spaces^18, 37, 40–42^. We found that none could execute on both the entire data set and the search space TagGraph considered in this comparison. Even by providing them with reduced number of spectra, search spaces, or both, TagGraph’s analysis speed was over an order of magnitude greater than the next fastest algorithm (Fig. 1c, Table 1).

**Table 1.** Search spaces considered by conventional and expanded database search algorithms.

| **Algorithm** | **Enzyme** | **Mods Considered** | **Mods/peptide** | **Protein Restriction** |
| --- | --- | --- | --- | --- |
| SEQUEST | LysC | methionine oxidation | three | **No** |
| Open search (SEQUEST) | LysC | Any mass between -500 and 500 Daltons | one (unlocalized) | **No** |
| PEAKS PTM | **None** | 435 previously known modifications (Unimod) | three | Yes |
| Byonic | LysC | Any mass between -40 and 100 Daltons | one | **No** |
| ModA | **None** | Any mass between -200 and 200 Daltons | **unlimited** | **No** |
| TagGraph | **None** | **Any mass** | **unlimited** | **No** |

### Effective error estimation for modified peptides: beyond target-decoy

Indexed string searches, as implemented by TagGraph, solve the long-standing conflict between search speed and search depth, but present a second challenge: estimating reliable false discovery rates (FDRs). The standard target-decoy estimation method we previously developed^32^ is unsuitable to TagGraph search results, since it loses discrimination accuracy as more peptides and PTMs are considered (**Supplementary Note 2**)^27, 43, 44^. Consequently, we developed a probabilistic validation strategy using a hierarchical Bayes Model optimized by expectation maximization (EM)^45^. Our robust model is universal, deducing the likelihood that any individual peptide-spectrum match is correctly interpreted, conditioned on fourteen quantitative and categorical attributes (**Supplementary Note 3**, Supplementary Fig. 3, Supplementary Fig. 4). Of these, half relate specifically to PTM-containing peptides, enabling discrimination between correctly and incorrectly interpreted spectra, regardless of the deduced peptides’ modification states.

We first evaluated TagGraph’s error model by comparing it to traditional target-decoy database search (SEQUEST), using the cell line dataset described in Fig. 1c. We found that EM-generated FDR estimates tended to be more conservative than those inferred from target-decoy searches (Fig. 1d), as expected for a model that discriminates between correct and incorrect PTM assignments. Furthermore, we found that the extent to which SEQUEST and TagGraph disagreed was consistent with the estimated 1% FDR threshold we applied to both (Supplementary Fig. 5a). For the majority of these disagreements however, TagGraph-generated peptide-spectrum matches were far more consistent with correct identifications based on protease specificity, algorithm-assigned scores, and ion assignment (Supplementary Fig. 5b, Supplementary Fig. 6).

To further evaluate TagGraph’s error model, we sought to measure how often modifications are miss-assigned to specific amino acid sites in the proteome. To accomplish this, we replaced all tyrosine residues with phenylalanines (mass difference of one oxygen, 15.9995 Da) in an altered human proteome sequence database (Fig. 1e). We reasoned that an accurate expanded search algorithm should return phenylalanine-containing peptides with an additional oxygen localized to converted phenylalanines; peptides containing converted phenylalanines without the oxygen addition are incorrect. This approach should therefore serve to benchmark modification assignments, which, similar to target-decoy’s use without modifications, delivers an expected known result, and could be generalized across multiple search engines.

We benchmarked four search methods against TagGraph with this validation tool. Each algorithm’s results were filtered based on target-decoy-based criteria (either the algorithm’s own implementation or a linear discriminant analysis^11^) or, for TagGraph, the hierarchical Bayes model. The proportion of peptide-spectrum matches containing unmodified phenylalanines at tyrosine positions was used to estimate the modification-specific FDR relative to the 1% predicted FDR. Due to their reliance on target-decoy based statistics (**Supplementary Note 2**), no algorithm besides TagGraph reliably discriminated phenylalanine-containing peptides: TagGraph’s error model was nearly an order of magnitude closer to the expected 1% than the next-best flexible search method (Fig. 1e), increasing sensitivity by more than four-fold (**Supplementary Table 1**) at 100 times the speed (Fig. 1c).

Even when searching the conventional human proteome sequence database, the four flexible search methods above produced ‘confident’ results with readily identifiable errors at severely underestimated FDR rates (**Supplementary Table 1, Supplementary Fig. 7**). In contrast, TagGraph doubled the number of unique peptide identifications relative to SEQUEST (Fig. 1f) by enabling accurate identification of peptides with any protease specificity and modification state. Once reconciled with the Unimod resource^10^, we found that unanticipated post-isolation modifications accounted for the majority of this increase (83%), followed by biologically-regulated modifications (12%) and those with no previous association (5%) (**Supplementary Table 1**). Our analysis of this cell line demonstrated TagGraph’s unique ability to sensitively characterize modified peptides with speeds and accuracies that are compatible with current, large-scale proteomic workflows.

### Unrestricted analysis of the human proteome reveals a broad modification landscape

To further investigate TagGraph’s utility for deep PTM characterization, we next applied our approach to a recently described draft human proteome^33^, approximately 150 times larger than our initial test data set. Due to their long computation times and underpowered validation techniques, performing this analysis with preexisting database search methods would not have been feasible. We interpreted 25 million tandem mass spectra derived from 30 adult and fetal tissues and over 2,000 raw data files^33^ with TagGraph. Once *de novo* sequencing (PEAKS ver. 7^46^) was complete, searching these data with TagGraph collectively took just six days on a single desktop computer. These data yielded over 1.1 million unique peptides, tripling the number originally reported using traditional database searching (Fig. 2a, **Supplementary Table 2**). This analysis identified proteins not found in the initial report, ranging from 100 (Adult CD8+ T Cells) to over 600 (Adult Gallbladder) additional proteins per tissue (Supplementary Fig. 8a, **Supplementary Table 3**). Several of these were supported by histological staining (Supplementary Fig. 8b).

**Figure 2.**
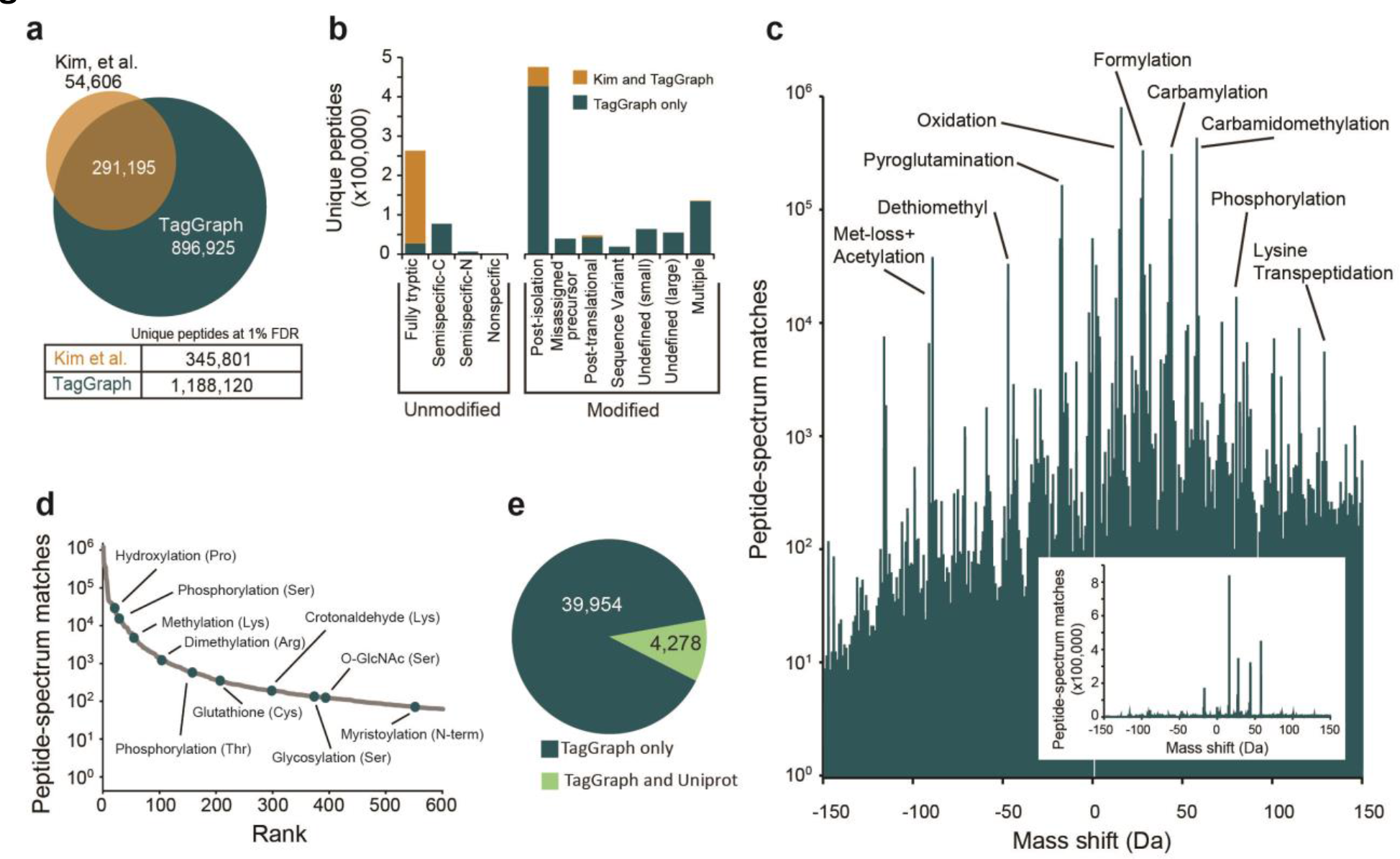
TagGraph extends deep proteome characterization to post-translational modifications. **a)** TagGraph confirmed the majority of identifications made by Kim et al.^33^ (**Supplementary Table 2**), but also expanded unique peptide identities from the human proteome dataset over three-fold relative to those originally reported. **b)** Categorical breakdown of unique peptide forms (distinguishing PTMs) identified by TagGraph. As expected, the majority of peptides identified by both TagGraph and Kim et al. correspond to tryptic peptides. Peptides identified by TagGraph but not Kim et al. primarily originated from non-tryptic peptides and peptides with unanticipated modifications. Post-isolation modifications comprised the most prevalent identification category in this dataset. **c)** Mass shifts (modified amino acid mass – unmodified amino acid mass) corresponding to all modifications identified by TagGraph from the human proteome dataset reveal a complex modification landscape. Numbers of identifications (peptide-spectrum matches) span six orders of magnitude. Despite the presence of several highly abundant post-isolation modifications (e.g., formylation), the depth of the proteomic profiling achieved in this dataset made it possible to characterize lower abundance post-translational modifications. Inset: modification frequencies without log transformation. **d)** Ranked relative abundances of 2,576 PTM-amino acid combinations, as estimated by the number of spectra bearing each from the human proteome dataset. Ten of these are highlighted; all modifications are represented in **Supplementary Table 4**. **e)** TagGraph analysis of the human proteome dataset identified 39,954 modification sites not present in Uniprot, 44,232 modification sites total. The overlap in the sites reported by TagGraph and Uniprot is highly significant (p-value < 1e-308, Fisher’s exact test).

**Table 2.** Top 10 Post-isolation, post-translational, and previously uncharacterized amino acid modifications identified from the Kim et al. dataset.

| Category<sup>a</sup> | Modification<sup>b</sup> | Specificity<sup>c</sup> | Mass Shift<sup>d</sup> | Unique Peptides<sup>e</sup> | Peptide-Spectrum Matches<sup>f</sup> | Sites<sup>g</sup> |
| --- | --- | --- | --- | --- | --- | --- |
| Post-isolation | Carbamylation | Peptide N-term | +43.01 | 63,302 | 254,382 | 70,207 |
|  | Formylation | Peptide N-term | +27.99 | 47,329 | 293,863 | 57,371 |
|  | Carbamidomethylation | Peptide N-term | +57.02 | 42,310 | 304,898 | 52,899 |
|  | Oxidation | Met | +15.99 | 42,193 | 784,001 | 41,115 |
|  | Deamidation | Asn | +0.98 | 21,064 | 252,713 | 22,609 |
|  | Acetaldehyde | Peptide N-term | +26.02 | 15,016 | 132,086 | 20,402 |
|  | Deamidation | Gln | +0.98 | 8,911 | 36,897 | 13,015 |
|  | Gln->pyroglutamate | Peptide N-term Gln | -17.03 | 8,372 | 123,277 | 7,966 |
|  | Carbamidomethylation | Lys | +57.02 | 7,575 | 40,422 | 12,355 |
|  | Carbamylation | Lys | +43.01 | 6,932 | 30,245 | 11,409 |
| Post translational | Phosphorylation | Ser | +79.97 | 3,466 | 15,798 | 3,729 |
|  | Met-loss + Acetyl | Protein N-term Met | -89.03 | 2,391 | 39,799 | 1,705 |
|  | Hydroxylation | Pro | +15.99 | 1,901 | 26,348 | 3,079 |
|  | Citrullination | Arg | +0.98 | 1,551 | 5,328 | 2,020 |
|  | GlyGly | Lys | +114.04 | 871 | 2,403 | 1,613 |
|  | Allysine | Lys | -1.03 | 861 | 2,801 | 1,670 |
|  | Hydroxylation | Lys | +15.99 | 766 | 3,316 | 1,090 |
|  | Cyano | Cys | -32.03 | 601 | 2,591 | 567 |
|  | Carboxylation | Glu | +43.98 | 467 | 726 | 913 |
|  | Acetylation | Lys | +42.01 | 455 | 1,323 | 1,196 |
| Previously undefined | Unknown | Peptide N-term | +12.00 | 3,834 | 14,743 | 3,692 |
|  | Unknown | Peptide N-term | +51.01 | 2,569 | 8,834 | 2,698 |
|  | Iron(III)<sup>h</sup> | Asp, Glu | +52.92 | 2,552 | 9,843 | 1,468 |
|  | carbamidomethyl and formyl on same residue<sup>h</sup> | Peptide N-term | +85.02 | 1,632 | 4,848 | 2,072 |
|  | disulfide bond<sup>h</sup> | Cys with nearby Cys | -116.06 | 1,529 | 9,788 | 1,840 |
|  | carbamidomethyl on C-terminus or Arg<sup>h</sup> | Peptide C-term Arg | +57.02 | 1,372 | 3,914 | 3,150 |
|  | Unknown | Peptide N-term | +83.04 | 1,280 | 4,632 | 1,948 |
|  | carbamidomethyl and pyro-glutamination on same residue<sup>h</sup> | Peptide N-term Glu | +39.01 | 1,099 | 3,622 | 908 |
|  | Unknown | Peptide N-term | +23.98 | 1,059 | 2,919 | 1,381 |
|  | reaction of N-terminal carbamidomethyl with internal Met<sup>h</sup> | Peptide N-term, co-occurs with dethiomethyl modification of internal Met | +105.02 | 957 | 4,907 | 991 |

a) Top ten modifications of the three indicated categories are shown, ordered by the number of unique peptides identified in the Kim *et al.* human proteome dataset. Categories assigned based on the likely modification identity, as determined by TagGraph.
b) Modification identities assigned by TagGraph, based on observed mass shifts, modification specificity, and evidence in the Unimod resource.
c) Specificity determined from the sites within modified peptides to which observed mass shifts were assigned.
d) Mass shift measured from the difference between an observed amino acid residue's monoisotopic mass and the expected value. Negative values indicate a net mass loss.
e) Number of unique peptide sequences bearing the annotated modification. Does not include peptide sequences identified with multiple modifications.
f) Total number of spectra in which indicated modification was confidently identified.
g) Total number of distinct amino acid residue sites bearing indicated modification.
h) Hypothetical identity of mass shift

As with our cell line analysis (Fig. 1), TagGraph predominantly rescued peptides bearing at least one modification that was not considered in the original search (Fig. 2b). A small number of post-isolation modifications (methionine oxidation; N-terminal carbamylation, carbamidomethylation, and formylation) collectively accounted for 38% of modified spectra (Fig. 2c, Table 2), consistent with previous findings^17, 18, 37, 47^. TagGraph rescued other commonly disregarded peptide classes, including semi-specific and non-specific trypsin cleavage, and mis-assigned monoisotopic precursor masses (Fig. 2b).

In comparison to the handful of abundant yet biologically irrelevant post-isolation modifications, this extremely deep proteome analysis revealed a much wider array of lesser-abundant PTMs (Fig. 2c-d, **Supplementary Table 4**). For example, we found N-terminal myristoylation, lysine hydroxylation, and arginine dimethylation hundreds to thousands of times in the proteome without requiring the kind of targeted, sample-intensive enrichment procedures that have previously been essential to PTM analysis. This study confirmed 4,278 modifications previously reported in the Uniprot proteomics resource, while extending it by an additional 39,954 (Fig. 2e, **Supplementary Table 5**). Comparing MS/MS spectra from this human proteome dataset to spectra derived from synthetic peptides (**Supplementary Fig. 9**) served to validate several unexpected, yet confidently identified peptides.

Many PTMs act as reversible switches on protein function. Their enzymatic addition and removal regulates signaling networks, protein binding, and other cellular processes^1, 3^. Although more than 90% of TagGraph-identified PTMs were previously unreported, we found several PTM-flanking sequence motifs^14, 48, 49^ enriched in this dataset (e.g. proline-directed phosphorylation^11, 50^ and glycine-directed arginine methylation^51^), supporting their validity (Fig. 3a, Supplementary Fig. 10). Furthermore, we identified over 200 gene ontologies^52^ that were significantly enriched among proteins bearing 22 noteworthy PTMs, giving additional support to their validity and functional significance (Fig. 3b, Supplementary Fig. 11, **Supplementary Table 6**). This unbiased analysis confirmed biological processes known to be regulated by multiple PTMs (e.g., acetyl Lys, methyl Lys, phosphorylated Ser regulating chromatin function^53^). Other processes, such as the cell cycle, were associated with a much more restricted set of PTMs (phosphorylated Ser)^54^.

**Figure 3.**
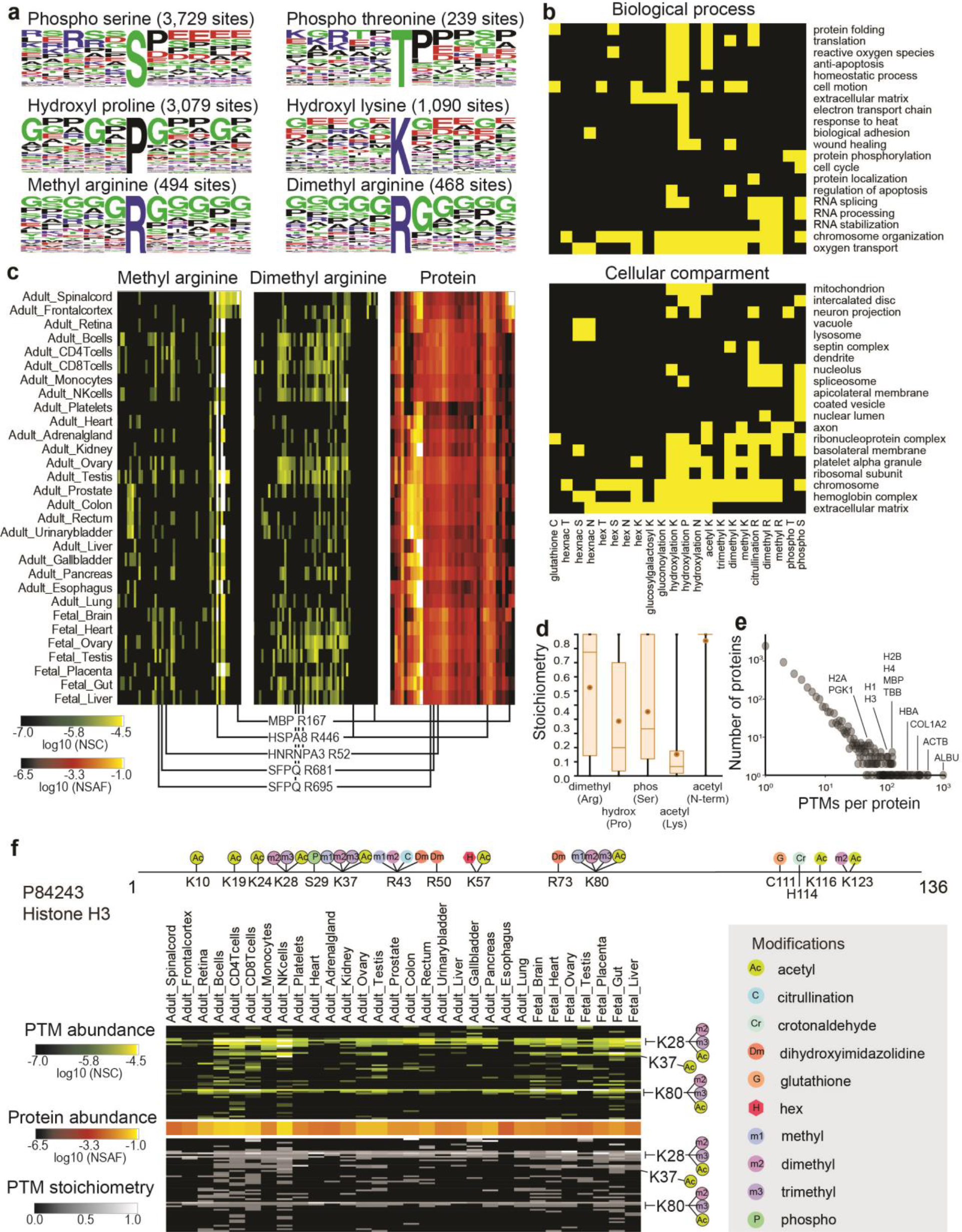
TagGraph reveals insights into PTM dynamics, function, and regulation. **a)** Sequence logos corresponding with select TagGraph-identified PTMs, as generated by WebLogo^87^. Amino acids flanking the indicated PTMs were evaluated with the Motif-X^48^ algorithm, confirming previously characterized motifs while positing new ones (Supplementary Fig. 10). **b)** Significantly enriched gene ontologies associated with prevalent post-translational modifications (yellow, 1% FDR (Benjamini-Hochberg corrected)). Ontologies and significances were assigned with the DAVID web tool^52^. A full list of all enriched ontologies is available in **Supplemental Table 6**. Ontologies significantly enriched among post-isolation modifications were excluded to correct for abundance-based biases in PTM detection (Supplementary Fig. 11). **c)** Arginine methylation and dimethylation distribution across proteins and tissues. The 64-most abundant monomethylated or dimethylated Arg sites from the entire data set are displayed across the y-axis, along with corresponding protein expression levels (49 proteins). Three modification sites on HNRNPA3 and SFPQ are highlighted for their distinct arginine monomethylation and dimethylation patterns across the tissues, despite demonstrating near uniform protein levels. Methyl modifications on MBP and HSPA8 are highlighted for their tissue specificity and ubiquity (respectively). Proteins were ordered by hierarchical clustering. PTMs were arranged to match their substrate proteins. All methylation sites are reported in **Supplementary Table 7**. **d)** Stoichiometry distributions vary for different PTMs, giving insight into their regulation and function. Box and whisker plots indicate the average (circle) and median (horizontal bar) values, 25^th^ quartile and 75^th^ quartile (box), and minimum and maximum (whiskers). **e)** Several proteins were found to be heavily modified in this data set. Histogram shows the number of proteins identified with the indicated number of distinct PTMs (site and modification). Of note, 921 distinct PTM sites were identified for human serum albumin. **f)** TagGraph identified both known and novel PTM sites on Histone H3 (**Supplementary Table 9**); a selection of the more abundant PTMs are shown. Site positions are numbered including the initiating methionine, as is the convention in the Uniprot protein database. PTMs circled in black are present in Uniprot. More detailed Histone PTM maps are presented in **Supplementary Table 9**.

We found that most PTMs were enriched in multiple biological process or cellular compartment were also implicated in multiple others. For example, reversible arginine methylation dynamically regulates proteins involved in RNA splicing and stabilization^50^, as confirmed by our ontology analysis (Fig. 3b). We observed a relative increase in the mono- and di-methylation site abundances on RNA splicing proteins such as HNRNPA3 and SFPQ in reproductive tissues and lymphocytes (Fig. 3c, **Supplementary Table 7**), suggesting that these modifications have specific roles in these contexts. Chemically similar modifications like arginine mono- and dimethylation showed stark contrasts: heat shock proteins were highly and consistently methylated in this dataset, but were not readily identified in dimethylated states (Fig. 3c, **Supplementary Table 7**).

### Quantifying PTM abundance and stoichiometry without requiring biochemical enrichment

We found that many PTMs’ abundances across the 30 tissues examined here mirrored those of the protein on which they were found, as exemplified by MBP R167 and HSPA8 R446 methylation (Fig. 3c). This degree of congruity suggests consistent stoichiometry across tissues. Conversely, other PTMs, including SFPQ R681 and R695 mono- and dimethylation, were largely restricted to specific tissues, despite the proteins’ uniform expression across the entire dataset (Fig. 3c). Since PTMs and their host proteins can be simultaneously quantified by mass spectrometry (e.g., Fig. 3c), we accordingly estimated each PTM’s stoichiometry (**Supplementary Methods, Supplementary Table 8**). This notion contrasts with previous PTM stoichiometry assays which required metabolic labeling^55, 56^, or enzymatic removal of a single target PTM class^56^. While such experimental interventions may estimate stoichiometries for a single PTM class (e.g., phosphorylation), it is difficult to use them to compare multiple, overlapping PTMs. Considering that a PTM’s stoichiometry can have important implications for its substrate protein’s activity and function^57^, deeply sequenced proteome datasets like this stand to illuminate a wide range of protein regulation.

In support of our flexible stoichiometry estimation approach, we found that protein N-terminal acetylation demonstrated the most consistently high stoichiometry (95.5%; stdev = 16.7%, Fig. 3d). This is expected, considering the broad and irreversible acetyl group addition, co-translationally catalyzed by N-terminal acetyltransferases^58^. Conversely, we found that lysine acetylation demonstrated consistently low and variable stoichiometry (15.2%; stdev = 22.7%, Fig. 3d), consistent with its heterogeneous representation on histone proteins^59^, and its possible non-enzymatic origins on abundant cytosolic and mitochondrial proteins^60, 61^. Over the entire dataset, we found that neither PTM abundance nor stoichiometry correlated with substrate protein abundance (Supplementary Fig. 12), supporting the complementary use of both measurements in proteome characterization.

### TagGraph simultaneously characterizes multiple PTM types on highly modified proteins

TagGraph identified multiple PTMs that intersect on individual proteins, and on individual residues (e.g., SFPQ R681, R695) (Fig 3c, **Supplementary Table 7**). Extreme examples include Albumin (921 PTMs) and actin (514 PTMs) (Fig. 3e). Histones are also well understood to undergo extensive and combinatorial modifications to encode epigenetic information^1^.

However, deciphering these modifications has required individual histone isoform^62^ or specific modification^63^ enrichment. Using TagGraph, we identified 277 PTMs across the major histone proteins, 132 of which were not previously reported (**Supplementary Table 9**). While we found modifications such as K28 dimethylation and K80 methylation on Histone H3 were both abundant and ubiquitous across the tissues examined here (Fig. 3f), we note several tissue-specific PTM combinations such as a 25-fold higher abundance of Histone H4 R56 dimethylation in fetal than adult tissues. Twenty-six PTMs, comprised of eleven PTM types, showed similarly higher abundance in fetal tissues, suggesting specific roles in developmental contexts (**Supplementary Table 9**). Our unbiased evaluation of these modifications, performed in conjunction with the rest of the proteome and without targeted enrichment techniques, opens new avenues to exploring tissue-specific epigenetic control.

### Enrichment-free PTM discovery identifies new roles for protein hydroxylation

We found that hydroxylation of prolines, tyrosines and lysines comprised a large (16%) proportion of newly identified histone PTMs (**Supplementary Table 9**), yet only hydroxylated tyrosine was previously described^64^. This coincides with our broader observation that several modification classes remain uncharted across the human proteome, despite being highly prevalent. Proline hydroxylation, for example, is the most abundant modification in the human body^65^, yet just 171 sites have been recorded^66^. Unlike more widely studied modifications, no enrichment tools exist to facilitate targeted hydroxylation analysis. Furthermore, of 11 amino acids capable of becoming hydroxylated^10^, four (Met, Trp, Phe, His) are often hydroxylated by standard proteomics sample preparation protocols. Thus, true post-translational proline hydroxylation must be distinguished from mis-localized artifacts^67^. Armed with TagGraph’s modification-focused error model, we confidently identified and localized 18-fold more hydroxyl proline residues than were previously known in humans (Table 2).

Proline hydroxylation is best understood in the context of collagen proteins, as it is essential to their role in maintaining extracellular matrix stability^65^. Despite hydroxyl proline comprising over 13% of mammalian collagen by weight^68^, only 128 sites across all collagens were previously assigned in humans (75% of all charted hydroxyl prolines in the human proteome). TagGraph identified 166 proline hydroxylation sites on COL1A2 alone, just three of which were previously described, identified by Edman degradation^69^(Fig 4a). While most proline hydroxylation sites were highly represented across most solid tissues examined here (e.g., P330, P642), several displayed tissue-specific abundance (e.g., P408, restricted to colon, bladder, liver, gallbladder, and pancreas) (Fig. 4b). TagGraph identified 25 other types of PTMs from this single protein, suggesting multiple routes by which PTMs cooperatively regulate collagen structure and function.

**Figure 4.**
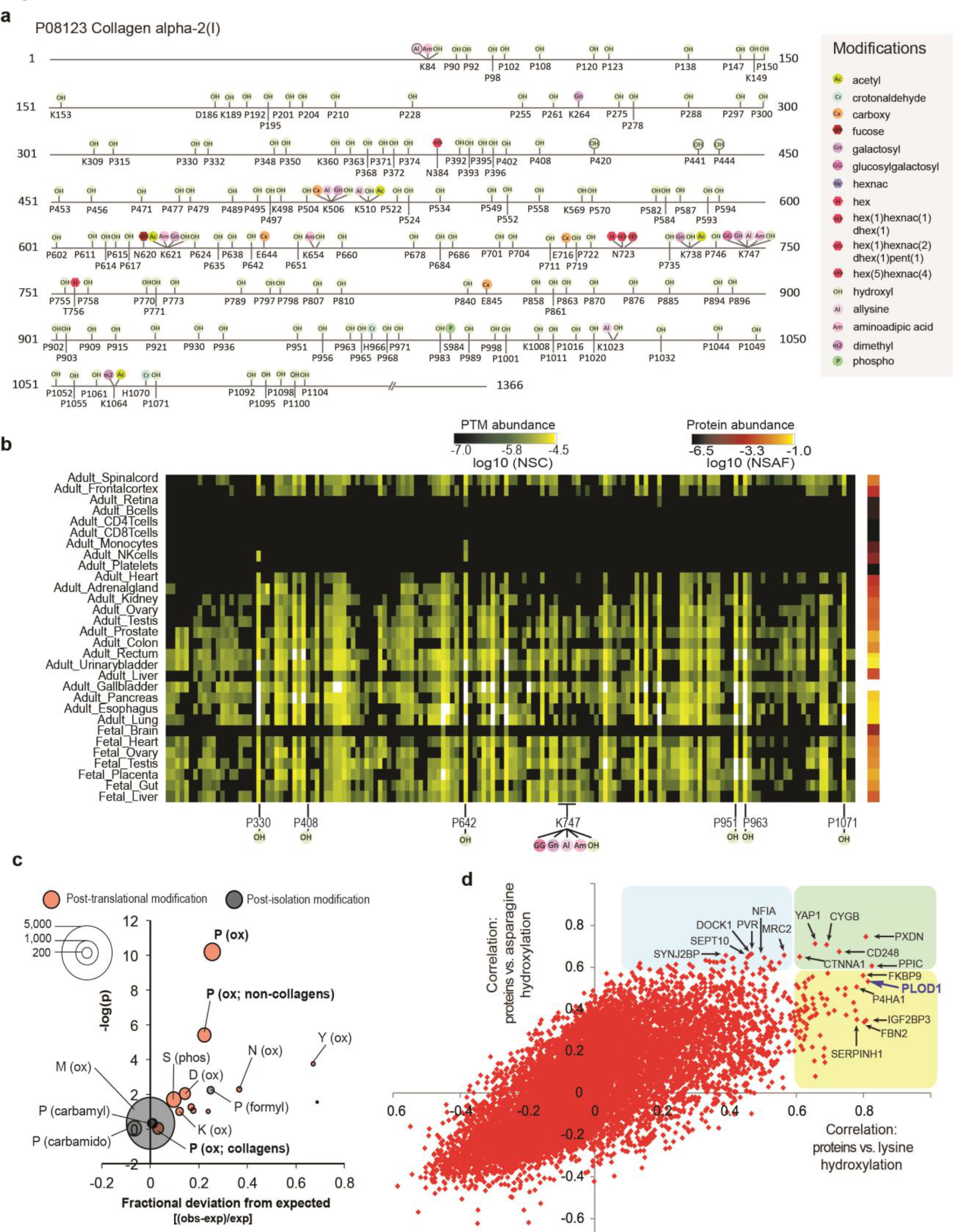
Characterization of hydroxylation, an un-enrichable PTM, enabled by TagGraph. **a)** TagGraph extensively expanded known hydroxylation sites across the human proteome; a selection of the most abundant PTMs on COL1A2 are shown as an example. TagGraph expanded proline hydroxylation from three previously known sites (P420, P441, P444)^69^ to 166 while identifying 25 other types of PTMs on this protein (**Supplementary Table 5**). **b)** PTMs identified in (a) were found to vary in abundance across tissues. Many hydroxylations displayed uniform abundance across solid tissues (i.e., P330, P642), whereas others displayed tissue-specific abundance variations (P408). **c)** Comparison between modification and cancer mutation sites (COSMIC). The size of each bubble indicates the number TagGraph-identified modification sites that were also found to be mutated in sequenced tumors. The expected value and significance (Fisher’s exact test) of this overlap was determined from the background of all peptides confidently identified by TagGraph (**Methods**). Proline hydroxylation sites significantly overlapped with mutation sites both overall (p <6e-11) and when restricted to non-collagen domain containing proteins (p <4e-6), suggesting that mutating these sites’ PTM capacity plays a role in cancer pathogenesis. **d)** Correlations between protein abundance and total PTM profiles across tissues (Supplementary Fig. 13) suggest candidate regulatory enzymes and functional associations. Proteins that highly correlated with lysine hydroxylation (x-axis), asparagine hydroxylation (y-axis) or both are highlighted (yellow, blue, or green, respectively). PLOD1 (in purple), the enzyme responsible for lysine hydroxylation in collagen emerged among the proteins most correlated with this modification. Protein expression levels were correlated with PTM stoichiometry across all tissues (**Methods**).

Although hydroxyl proline’s role in maintaining collagen structure is well understood^65^, its prevalence and roles on other proteins has remained sparse^70–72^. Just 26 proteins were previously reported to bear hydroxyl proline modifications besides collagens and collagen domain-containing proteins^66^, and thus, it has widely been considered a relatively specialized PTM. Our analysis extends known proline hydroxylation by nearly 3,000 sites spanning nearly 1,000 substrate proteins (**Supplementary Table 5**). These proteins were significantly enriched for 113 biological processes (**Supplementary Table 6**). Thus, proline hydroxylation likely shapes a diverse range of cellular processes beyond matrix homeostasis (Fig. 3b).

Noting that tumors also exploit multiple cellular processes during oncogenesis, we hypothesized that proline hydroxylation could play a role in cancer. Significant associations were previously shown between specific phosphorylation sites and cancer-associated mutations^8^. Taking a similar approach, we examined whether proline hydroxylation significantly intersected with missense somatic cancer mutations catalogued in the COSMIC database^73^. We found that hydroxylated prolines were 25% more likely to be associated with cancer mutations than expected (p<6e-11, Fisher’s exact test, Fig. 4c). Enrichment persisted even after excluding collagen domain-containing proteins (22%, p<4e-6, Fig. 4c). Methionine oxidation, a common post-isolation modification, was not enriched (p=0.49), nor were other post-isolation proline modifications (Fig. 4c). Similar to phosphorylation, further study of hydroxylation could substantially increase insight into cancer pathogenesis and reveal new therapeutic targets. The identification of mutated hydroxylation sites on proteins which are hubs of post-translational signaling (i.e., 20 such sites on histones) further supports this hypothesis (**Supplementary Table 10**).

### Identifying candidate PTM interactors and regulators

As with proline hydroxylation, TagGraph significantly expanded the number of known sites for lysine hydroxylation and asparagine hydroxylation by 18-fold and 14-fold, respectively. To further elucidate protein-PTM interactions beyond ontological groupings, we reasoned that PTMs should co-occur with the specific proteins with which they interact. We tested this notion by screening all PTM and protein quantifications for significant correlations across the 30 tissues examined here. We found several protein-PTM correlations that confirmed known functional associations (Fig. 4d, Supplementary Fig. 13). Generally, we found that proteins that were highly correlated with specific modifications did not bear those modifications themselves (Supplementary Fig. 13a). However, they tended to be enriched for the same functional ontologies as the PTMs’ substrates (Supplementary Fig. 13 c). Such highly correlated proteins are functionally associated with these PTMs and may be candidate PTM-altering enzymes or indirect regulators.

We found over 70 proteins with abundances that correlated highly with lysine hydroxylation abundance across all tissues (Fig. 4d). Many of these proteins, such as PXDN and CYGB have known roles in oxygen transport or oxidoreductase activity (Supplementary Fig. 13c), supporting their role in regulating hydroxylation PTMs. Of note, one enzyme known to catalyze this PTM, PLOD1^74^, was the second most highly correlated (Fig. 4d).

Surprisingly, many proteins that correlated with lysine hydroxylation also correlated with asparagine hydroxylation (Fig. 4d), despite no previous evidence linking these PTMs to the same biological context. The primary substrates for lysine hydroxylation are collagens, which are a major component of the extracellular matrix (ECM)^75^. Though asparagine hydroxylation was not previously characterized in the ECM, TagGraph revealed 45 novel sites on Fibrillin-1 and Fibrillin-2 (**Supplementary Table 5**), both of which are ECM constituents. As opposed to specific correlates (such as PLOD1 for lysine hydroxylation), proteins that correlate highly with both PTMs may function as general positive regulators of ECM homeostasis through hydroxylation (Fig. 4d). It is plausible that asparagine hydroxylation might stabilize fibrillins, analogous to the function of lysine and proline hydroxylation in stabilizing collagen fibrils^65^ and asparagine hydroxylation in stabilizing ankyrins^76^.

## Discussion

Characterizing the identity, abundance, and function of post-translational modifications (PTMs) is arguably the single-most important contribution mass spectrometry-based proteomics can make to cell biology^16^. However, computational and experimental limitations have reinforced a myopic view of a diverse and dynamic PTM landscape. TagGraph overcomes these obstacles with two major innovations, making it possible to identify essentially any modified peptide sequence from high-quality tandem mass spectra.

First, we circumvent the slow task of performing thousands or tens of thousands of peptide-spectrum comparisons for each observed tandem mass spectrum. Instead, we devised an extremely rapid string matching approach that produces only a handful of candidate sequence matches that are then scored against the observed spectrum. A graph-based reconciliation algorithm lets TagGraph consider any combination of modifications to a peptide. Importantly, it performs this task at speeds commensurate with conventional database search algorithms. Just as similar string matching algorithms revolutionized next-generation DNA sequencing^31^, we expect this capability will become increasingly important as the latest generation of high-resolution and high-volume mass spectrometers^77, 78^ become more widely available.

Second, we optimized a probabilistic model that simultaneously evaluates the likelihood that a peptide’s sequence and any modifications it bears are correct. This component of our approach was essential, since the standard “target-decoy” error estimation method we previously developed is inherently blind to amino acid modifications^67, 79^. We demonstrate the accuracy of our error estimations using synthetic peptide validation, direct comparison to target-decoy, and re-identification of known sites, motifs, protein-PTM relationships, and functional roles for various PTMs. We expect the ease with which the model can be modified will offer superior FDR estimation to target-decoy in several other specific applications^80, 81^.

Our analysis of an unprecedentedly rich PTM landscape could only be accomplished through these two advances. This kind of high-throughput, unbiased PTM discovery is compatible with any proteomics experiment using high-resolution tandem-mass spectrometry. We demonstrate this ability in several ways, including a focused analysis on proline hydroxylation, a pervasive PTM previously only known to occur on a small number of proteins. We expand the number of known sites by 18-fold, linking proline hydroxylation to a much wider array of biological contexts than originally thought, and demonstrate its significant association with somatic cancer mutations. Furthermore, simultaneous identification of unmodified and modified peptides from the same dataset enables high throughput quantification of PTM stoichiometries. This metric holds great potential for illuminating functional relationships between PTMs, their substrates and candidate regulatory proteins.

In addition to enabling routine, enrichment-free PTM discovery and characterization, we envision many other applications for TagGraph. By searching MS/MS spectra in an enzyme-independent manner, for example, TagGraph could automatically detect endogenous peptides ^82^ and alternate start site utilization^83^. Furthermore, TagGraph’s speed holds tremendous advantages for refining gene predictions when applied in a proteogenomic context: by rapidly evaluating multiple gene assemblies in all six frames while permitting amino acid substitutions, insertions, and deletions it can validate translation products that would confound conventional database search methods. This capacity could have direct application to systems that have previously been intractable to large-scale proteome analysis such as the gut microbiome and other complex microbial communities. Finally, by learning essential experimental details (e.g., PTMs, enzymatic digestion, and mass accuracy) directly from input data, TagGraph can help standardize proteomic analyses. This capability stands to enable direct comparisons between data sets collected by multiple laboratories, thereby fostering the kind of large-scale collaborations that have transformed the genomics field.

## Acknowledgements

This work was supported by a Damon Runyon-Rachleff Innovation Award from the Damon Runyon Cancer Research Foundation (DRR-13-11; J.E.E), the W.M. Keck Foundation Medical Research Program (J.E.E.), the Bill and Melinda Gates foundation (J.E.E.), the Stanford Graduate Fund (A.D.), the Wallenberg Foundation (N.O.), and the National Science Foundation Graduate Research Fellowship (C.G.). Thanks to Ross Flockhart and Paul Khavari for supplying A375 cells for our pilot studies. Thanks to Aparna Badhuri for assistance with data visualization. We further acknowledge members of the Elias Lab, as well as David Dill, Paul Khavari, Parag Mallick, Tobias Meyer, Daria Mochley-Rosen, and Joanna Wysocka for helpful discussions during the preparation of this manuscript.

## Author contributions

A.D. designed and implemented all algorithms, designed and carried out all A375 mass spectrometry experiments, performed the human proteome analysis, and wrote the manuscript. J.E.E. designed algorithms and wrote the manuscript N.O. performed synthetic peptide spectra analysis. K.R. implemented TagGraph for web-based queries. C.G. wrote the manuscript. K.S. performed experimental validation studies and wrote the manuscript.

## SUPPLEMENTARY METHODS

### 1. Datasets

#### a. A375 data set

##### 1.a.i Sample processing

A375 melanoma cells (ATTC)^1^ were cultured in DMEM supplemented with 10% FCS and antibiotics. Cells were detached by trypsinization, pelleted, washed with PBS and flash frozen in liquid nitrogen. 5×10^7^ flash-frozen A375 cells were thawed on ice and lysed by tip sonication in Urea lysis buffer (8 M Urea, 100 mM NaCl, 50 mM Tris, 1 mM PMSF, 10 µM E-64, 100 nM bestatin, pH 8.2). The cell lysate was reduced (5 mM DTT, 55 °C, 30 min), alkylated (12.5 mM iodoacetamide, room temperature, 1 hr in the dark), and digested overnight with LysC at an enzyme:substrate ratio of 1:100 (37 °C). The resulting peptide mixture was desalted using C-18 Sep-Pak cartridges (Waters), dried using vacuum centrifugation, and resuspended in 10 mM ammonium formate, pH 10 prior to high pH reverse phase (HPRP) separation. HPRP was performed using an Agilent 1100 binary HPLC, delivering a gradient (0%-5% B over 10 min, 5%-35% B over 60 min, 35%-70% B over 15 min, 70% B for 10 min) across an Agilent C-18 Zorbax Extend column. Buffer A was 10 mM ammonium formate, pH 10 and buffer B was 10 mM ammonium formate, 90% acetonitrile, pH 10. Sixty one-minute fractions were collected and concatenated into twelve fractions as described previously ^2^.

##### 1.a.ii Mass Spectrometry

All HPRP fractions were desalted using C-18 Sep-pak cartridges (Waters), vacuum centrifuged, and resuspended in 5% ACN, 5% formic acid at approximately 1 ug/ul. One microgram of each fraction was analyzed by microcapillary liquid chromatography electrospray ionization tandem mass spectrometry (LC-MS/MS) on an LTQ Orbitrap Velos mass spectrometer (Thermo Fisher Scientific, San Jose, CA) equipped with an in-house built nanospray source, an Agilent 1200 Series binary HPLC pump, and a MicroAS autosampler (Thermo Fisher Scientific). Peptides were separated on a 125 um ID x 18 cm fused silica microcapillary column with an in-house pulled emitter tip with an internal diameter of approximately 5 um. The column was packed with ProntoSIL C_18_ AQ reversed phase resin (3 um particles, 200Å pore size; MAC-MOD, Chadds Ford, PA). Each sample was separated by applying a two-step gradient: 7% -25% buffer B over 2h; 25-45% B over 30 min. Buffer A was 0.1% formic acid, 2.5% ACN and buffer B was 0.1% formic acid, 97.5% ACN. The mass spectrometer was operated in a data dependent mode in which a full MS scan was acquired in the Orbitrap (AGC: 5×10^5^; resolution: 6×10^4^; m/z range: 360-1600; maximum ion injection time, 500 ms), followed by up to 10 HCD MS/MS spectra, collected from the most abundant ions from the full MS scan. MS/MS spectra were collected in the Orbitrap (AGC: 2×10^5^; resolution: 7.5×10^3^; minimum m/z: 100; maximum ion injection time, 1000 ms; isolation width: 2 Da; normalized collision energy: 30; default charge state: 2; activation time: 30 ms; dynamic exclusion time: 60 sec; exclude singly-charged ions and ions for which no charge-state could be determined).The mass calibration of the Orbitrap analyzer was maintained to deliver mass accuracies of ±5 ppm without an external calibrant. All raw mass spectrometry data were uploaded to the PRIDE repository^3^ and assigned the reference ID PXD######.

#### 1.b. Synthetic peptide confirmation data set

In total, 86 synthetic peptides (SpikeTides from JPT Peptide Technologies GmbH) validating various modifications, semi- or non-specific peptide assignments were evaluated. A final pool of all peptides dissolved in 0.1% formic acid was generated and the concentration of each individual peptide was roughly 250 fmol/µL. Two LC-MS/MS runs were performed and the auto-sampler injected 1 µl of the synthetic peptide pool. An ESI-Orbitrap Elite mass spectrometer (Thermo Electron, Bremen, Germany) interfaced with an Eksigent ekspert nanoLC 425 system (Eksigent technologies, Dublin, CA, USA) was used. Peptides were introduced into the mass spectrometer via a fused silica microcapillary column (100 µm inner diameter) ending in an in-house pulled needle tip. The columns were packed in-house to a length of 18 cm with a C18 reversed-phase resin (with Reprosil-Pur C18-AQ resin (3 µm Dr. Maisch, GmbH, Germany). For elution a two-step gradient of 4-25% buffer B (5 % DMSO, 0.2% formic acid and 94.8 % acetonitrile (v/v)) in buffer A (5 % DMSO, 0.2% formic acid in water (v/v)) over 60 min followed by a second phase of 25-45% buffer B over 20 min was used. The LTQ-Orbitrap was operated in data-dependent mode to automatically switch between Orbitrap-MS (from *m/z* 340 to 1600) and ten MS/MS acquisition. Each FT-MS scan was acquired at 60,000 FWHM nominal resolution settings while the MS/MS spectra were acquired using HCD and at a resolution of 15,000. Precursor ion charge state screening was enabled (charge state 1 rejected) and the normalized collision energy was set to 35%.

The resulting data were analyzed by TagGraph to match mass spectra with their best-matching synthetic peptide sequence. Synthetic-derived and experiment-derived mass spectra (e.g., from the draft human proteome data set) were only compared if spectra were assigned to the same peptide sequence in the same charge state. Of the 86 peptides synthesized, 75 were matched in this manner and used to validate TagGraph peptide-spectrum assignments (**Supplementary Fig. 9**). The mass tolerance used to match b- and y-ions was 0.1 Daltons.

#### 1.c. Draft human proteome data set

All RAW data and database search results from the draft human proteome data set ^4^ were downloaded from the PRIDE^5^ data repository using accession PXD000561.

### 2. *De novo* search engine comparisons

We compared the performance of three *de novo* sequencing algorithms, PEAKS 7 ^6^, PepNovo+ (ver. 3.1) ^7^, and pNovo (ver 1.1) ^8^ and assayed their ability to generate *mostly* correct sequence interpretations of high mass accuracy MS/MS spectra^9^. Each algorithm was used to search the A375 dataset, and the resulting peptide identifications were compared against high confidence peptide spectrum matches obtained from a SEQUEST search of the same dataset as previously described^9^(**3**, below). Static and differential modifications were set as for the SEQUEST search. Mass tolerance parameters were optimized to achieve maximum sequencing accuracy for each algorithm individually^9^. PEAKS was run with a precursor mass tolerance of 10 ppm and a fragment mass tolerance of 0.01 Da. PepNovo+ was run a 0.01 Dalton precursor mass tolerance and 0.05 Da mass tolerance on fragment ions. pNovo was run with a 6 ppm precursor mass tolerance and a 25 ppm fragment ion tolerance.

The accuracy of each *de novo* algorithm was assessed using the **sequence accuracy** metric (Supplementary Fig. 2a)^9^. For a given *de novo* peptide-spectrum match and its corresponding high confidence SEQUEST peptide-spectrum match, sequence accuracy represents the fraction of prefix residue masses^10^ present in the high confidence SEQUEST match which were also present in the *de novo* sequence^9^.

### 3. Database search engine comparisons

We assessed several expanded database search algorithms’ abilities to detect undefined modifications, without constraining protease specificity, using the A375 dataset. As a baseline, we searched all 168,391 MS/MS spectra in this dataset with SEQUEST ^11^ (ver. 28 rev 12) using an indexed sequence database comprised of the human proteome (Uniprot, downloaded 12/9/2014) ^12^ plus common contaminants. The database was concatenated with a reversed database for target-decoy FDR estimation. The SEQUEST search was conducted with LysC protease specificity, 50 ppm precursor ion mass tolerance, and 0.5 Da fragment ion mass tolerance. Cysteine carbamidomethylation (+57.021464 Da) was set as a static modification and methionine oxidation (+15.994915 Da) was set as a differential modification.

The A375 dataset was then searched using PEAKS PTM (ver. 7) ^13^, Byonic (ver. 2.5.6) ^14^, ModA (ver. 1.03) ^15^, and the Open search method using SEQUEST ^16^ – four strategies described as being able to either consider relatively large numbers of discrete amino acid modifications, or searching spectra with no *a priori* constraints on possible modifications. It was not possible to search the entire A375 dataset with any of the above algorithms using completely unconstrained parameters with respect to both modifications and protease specificity: either the algorithms would not execute, or they did not complete within a reasonable amount of time (5 days per RAW data file). Thus, CPU times were calculated using the most feasible parameters approximating such a search for each algorithm, and extrapolated from a limited subset of search results in cases where searching the entire dataset would be too computationally intensive. As such, the search times reported in Fig. 1d represent a substantial underestimate of the true time needed for each of these algorithms to analyze a sample in a manner equivalent to TagGraph, as described below.

For PEAKS PTM, the A375 dataset was first *de novo* sequenced using the following settings: 10 ppm precursor ion tolerance and 0.01 Da fragment ion tolerance, cysteine carbamidomethylation as a static modification, and methionine oxidation as a differential modification. The dataset was then analyzed with PEAKS PTM using the same modification and mass tolerances as for the *de novo* sequencing, LysC enzyme specificity allowing for nonspecific cleavage at both ends of the peptide, and additionally searching with all 485 modifications curated in PEAKS’s internal PTM database. For ModA, the A375 dataset was analyzed with the following settings: 0.05 Da precursor mass tolerance, 0.05 Da fragment ion tolerance, allowing one modification per peptide, no protease specificity, modification size between -200 Da and 200 Da. For Byonic, the dataset was analyzed using the following settings: 10 ppm precursor ion tolerance, 20 ppm fragment ion tolerance, LysC protease specificity, cysteine carbamidomethylation as a static modification, methionine oxidation as a differential modification, wildcard search enabled with a minimum mass of -200 Daltons and a maximum mass of 200 Daltons. For all three algorithms, the sequence database used was the same as for SEQUEST. PEAKS PTM and Byonic were allowed to use their own internal methods for creating decoy sequences, whereas ModA was given a concatenated forward/reversed database as input. The open search method was conducted using SEQUEST and the same indexed database as above, with a 500 Da tolerance on the precursor ion mass and a 0.1 Da mass tolerance on fragment ion masses.

For the search time comparison between all algorithms (Fig. 1d), CPU times were calculated as the sum of the CPU time over all processes spawned by each database search algorithm to analyze the data. Due to computational constraints, it was not possible to run Byonic with a semi specific or nonspecific enzyme specificity, or to search the entire A375 dataset with either Byonic or ModA. Thus, we conducted the Byonic search with full LysC specificity, and analyzed only a single fraction of the A375 dataset for both Byonic and ModA. The estimated CPU time over the entire dataset was extrapolated by multiplying the CPU time recorded from the analysis of a single HPRP fraction by the ratio of the total MS/MS spectra in the dataset (168,391) divided by the number of MS/MS spectra in the single fraction (16,613).

### 4. TagGraph Parameters

TagGraph was used to analyze both the A375 dataset and the draft human proteome data set. In both cases, all available MS/MS were first *de novo* sequenced using PEAKS. The resulting peptide sequences and raw mass spectra (mzXML-formatted ^17^) were given as input to TagGraph.

For the A375 dataset, PEAKS was run as described above. For the draft Human Proteome data set, PEAKS was run with a 10 ppm precursor mass tolerance, 0.05 Dalton fragment ion tolerance, cysteine carbamidomethylation as a static modification, and methionine oxidation as a differential modification to maximize sequence accuracy.

The *de novo* sequencing results were searched with TagGraph against the human proteome (Uniprot, downloaded 12/9/2014) plus common contaminants. The database was concatenated with reversed decoy sequences only for searches of the A375 dataset. This enabled direct comparisons of FDR estimates derived from TagGraph with those derived from target-decoy searching, and fair comparison of the CPU time of TagGraph with the other database search algorithms. For the draft human proteome data set, only the forward database was searched as FDR estimates were derived using the hierarchical Bayes scoring model. For both datasets, mass tolerances were set to 10 ppm precursor ion tolerance and 0.1 Dalton fragment ion tolerance. Enzyme specificity was set to LysC for the A375 dataset and Trypsin for the human proteome data set. Although enzyme specificity was considered as a scoring attribute in the hierarchical Bayes model, TagGraph is able to return high-confidence semi specific and nonspecific peptide-spectrum matches regardless of the input enzyme specificity.

High confidence results were retrieved at a 1% FDR for the A375 dataset by ranking all returned peptide-spectrum matches according to their probabilities P(D|+) from highest to lowest, then adding matches in order of decreasing rank to the set of high confidence results until the expected FDR equaled 1% (**Supplementary Note 3**, Equation 2). The human proteome data set was evaluated in a similar manner, considering each experiment (e.g., gel fractionation of adult heart tissue) individually rather than for the entire dataset as a whole.

### 5. PTM FDR estimation with amino acid-substituted proteome

Target-decoy based error estimation accuracy declines when applied to peptide modifications and other large search spaces ^18^ (**Supplementary Note 2**). Despite this, all previously described unrestricted search algorithms rely on target-decoy to delineate sets of confidently identified spectra. To assess the degree to which predicted FDRs produced by these algorithms deviate from the actual FDR, we employed a modified human proteome sequence database in which every tyrosine residue was replaced by a phenylalanine. The mass difference between these residues (15.994915 Da) corresponds with an oxygen atom, and is a frequently observed modification on several residues (e.g., methionine), while distinguishing other unmodified residues (e.g., alanine and serine). Thus, we reasoned that search engines capable of accurate PTM assignment and discrimination should search the tyrosine-substituted database and return phenylalanine-containing peptides modified by an oxygen only on those phenylalanines that were previously tyrosines. They should be able to discriminate these identifications from erroneous ones in which oxidation modifications were assigned to unaltered residues. We analyzed the A375 dataset against this modified sequence database using SEQUEST, PEAKS PTM, Byonic, and ModA. The results from each algorithm were then filtered to a 1% predicted FDR using target-decoy based statistics. Byonic and PEAKS PTM were allowed to use their own internal target-decoy based filtering algorithms. Search results provided by SEQUEST and ModA were filtered using a linear discriminant method ^19^. The FDR for each set of search results was calculated as the proportion of peptide-spectrum matches containing phenylalanines at tyrosine positions which were not annotated with a phenylalanine to tyrosine modification (+15.9995 Da) or a modification of equivalent mass.

The SEQUEST search was conducted with phenylalanine to tyrosine, methionine hydroxylation, proline hydroxylation, lysine hydroxylation, and asparagine hydroxylation as differential modifications. Aside from this change, all searches were conducted with the same parameters and data as described in section 3 above. Open search was not considered in this comparison as it does not provide amino acid localizations for its predicted modification masses.

The A375 dataset was also analyzed with TagGraph against the modified human sequence database. The parameters used were identical to those described in section 4 above. Results were filtered to a 1% predicted FDR using the hierarchical Bayes model and the actual FDR was calculated as described above.

### 6. Abundance calculations

#### 6a. Protein abundance calculation

Protein abundances were calculated using the distributed normalized spectral abundance factor (NSAF) method ^20^. Briefly, the number of spectral counts originating from peptides that uniquely map to single proteins were summed over all proteins identified in an experiment. Spectral counts recorded from peptides that map to multiple proteins were distributed across all such proteins according to the proportion of spectral counts assigned to them from uniquely mapped peptides. Finally, summed spectral counts for each protein were normalized by protein length, and the sum of all protein abundances for each experimental dataset was normalized to one.

Protein abundances per tissue were calculated as the average of the individual NSAF for that protein over all experiments performed on that tissue.

#### 6b. Site abundance and stoichiometry calculations

To compare modification sites between tissues, we quantified the abundance of sites using two methods: normalized spectral counts (NSC) and estimated stoichiometry. For both methods, we first generated a catalog of all confident peptide identifications that span a given modified amino acid position of a protein, regardless of modification state. The total spectral counts corresponding to all peptides containing the amino acid position (S_T) and just those corresponding to peptides containing the exact modification on the site of interest (S_m) were calculated for each experimental dataset.

The normalized spectral count of a modification site is calculated as S_m divided by the number of confidently identified peptide-spectrum matches in the experimental dataset. The stoichiometry of the modification is calculated as S_m divided by S_T. Modification site abundances (stoichiometry or NSC) per tissue were calculated as the average of the site abundances over all experiments performed on that tissue. Due to inherent difficulties in accurately reporting very low abundances with spectral counting ^21^, experiments in which no peptides overlapping the site of modification were detected were not included in the average.

Thus, the sum of stoichiometries of all modifications at a particular site in a particular tissue may not be normalized. Finally, the abundance, stoichiometry or normalized spectral count of a modification site was set to zero for a particular tissue if the corresponding protein NSAF was zero in that tissue.

### 7. Gene ontology analysis

Gene ontology analysis was conducted using the DAVID web portal ^22^. For each post-translational modification of interest, proteins bearing that modification were compiled and input as a gene list. The background list used was the Uniprot human proteome. The resulting gene ontologies were downloaded and a global FDR threshold (Benjamini-Hochberg) of 1% was used as a threshold for determining significantly enriched ontologies.

We observed that many ontologies were broadly enriched across all post-translational modifications and hypothesized that these were simply associated with highly abundant proteins and did not reflect true post-translational modification properties. As a control, we applied the above enrichment analysis to fifteen post-isolation modifications and observed many ontologies that were significantly enriched for all post-isolation modifications considered (Supplementary Fig. 11). These ontologies were excluded from the set of enriched ontologies in the post-translational modification analysis (Fig. 3b, **Supplementary Table 6**).

### 8. COSMIC dataset comparison

A database of cancer mutations was downloaded via FTP from the COSMIC website^23^. The mutation list was then filtered to keep only missense mutations. To guard against slight variations in protein sequence between COSMIC and Uniprot, mutations for which the amino acid residue at the denoted position in the Uniprot protein sequence did not match the non-mutated amino acid identity in the corresponding COSMIC entry were discarded.

To guard against biases in background amino acid distributions, overlap statistics were only calculated for proteins on which both cancer mutations and the PTM of interest was detected and only against the background of peptides confidently identified by TagGraph in the human proteome dataset. Using proline hydroxylation as an example, the number of prolines, number of hydroxylation prolines, number of mutated prolines, and number of mutated and hydroxylated prolines were counted only on peptides confidently identified by TagGraph and on proteins containing both cancer mutations and proline hydroxylation. This overlap was then tested for significance via Fisher’s exact test. This analysis was carried out analogously for other types of hydroxylations (lysine, asparagine, methionine, etc.).

### 9. Protein-PTM correlation analysis

Reasoning that many modifications’ abundances and stoichiometries will depend on specific protein-modifying enzymes, we sought to discover functional relationships between post-translational modifications and proteins. We identified highly correlated subsets of modifications and proteins by comparing their abundances across the tissues examined here. Modification site and protein lists were first filtered to include only those identified from at least three tissues. For a particular post-translational modification of interest (e.g., Lysine hydroxylation), the abundance of the modification was averaged across all measured sites from all proteins within each tissue, forming a vector representing the abundance of the modification across all tissues. Similarly, for all identified proteins, the calculated NSAF was used to form an abundance vector of that protein’s expression across all tissues. We next determined the Pearson correlation coefficient between all modification and all protein vectors computed and filtered as described above. The proteins with the largest magnitude correlations (positive or negative) were then considered as candidates having a functional relationship with a modification of interest.

Modification abundance vectors were calculated using both modification stoichiometries and modification-normalized spectral counts. Both types of quantification were used in the correlation analysis, often yielding different results (**Supplementary Fig. 13a**). However, in both cases, our analysis revealed previously described associations between proteins and post-translational modifications (e.g., arginine methylation and RNA splicing proteins), supporting the validity of this analysis (**Supplementary Fig. 13c**).

### 10. Code availability

The TagGraph algorithm is currently available via a web interface at http://kronos.stanford.edu/TAG_GRAPH/. The source code is available through github at http://github.com/adevabhaktuni/XXXXXXXXX

## SUPPLEMENTARY FIGURES

**Supplementary Fig. 1.**
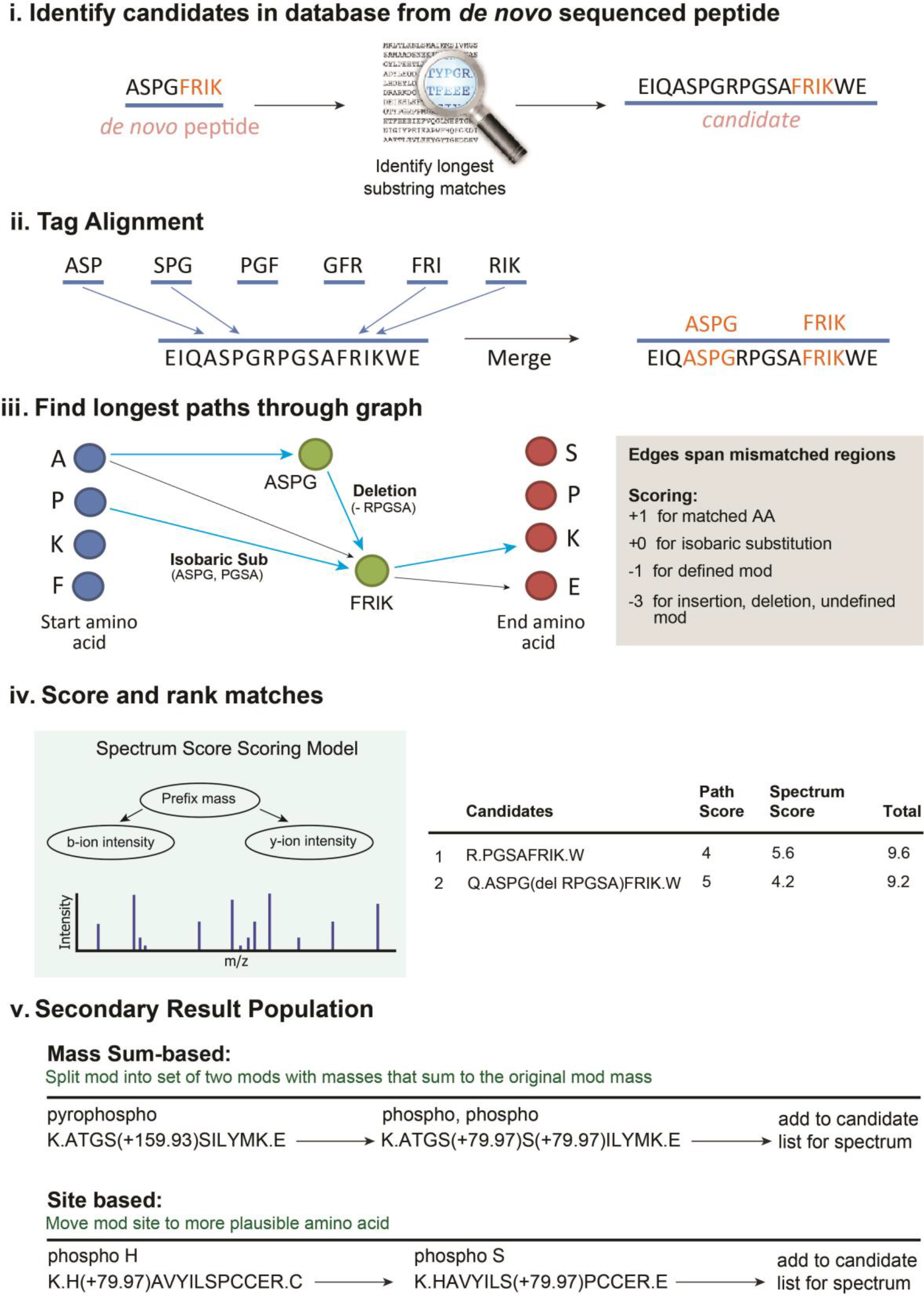
TagGraph algorithm workflow overview. TagGraph employs a five-step procedure as depicted below, and detailed in **Supplementary Note 1**: **(i)** *De novo* sequences are used to query an indexed sequence database. All candidate database entries containing a maximum-length substring in common with the *de novo* sequence are retrieved. **(ii)** The *de novo* sequence is compared against each database-derived candidate match. Continuous amino acid substrings of length >2 that are identical between the query and database candidate are identified as putative tags. **(iii)** Candidate matches (defined as a peptide plus the set of its assigned modifications) are retrieved using a longest path algorithm on a directed acyclic graph. Sequence tags defined in (ii) above are represented as nodes in the graph and modifications as edges. Paths are drawn from start positions on the database peptide to end positions through nodes and edges. **(iv)** Candidate matches over all database peptides are collected and scored against the MS/MS spectrum using a probabilistic scoring model. **(v)** After all *de novo* sequences are analyzed, additional candidate modification annotations are created for select spectra if they are likely to be correct based on global dataset modification abundances.

**Supplementary Fig. 2.**
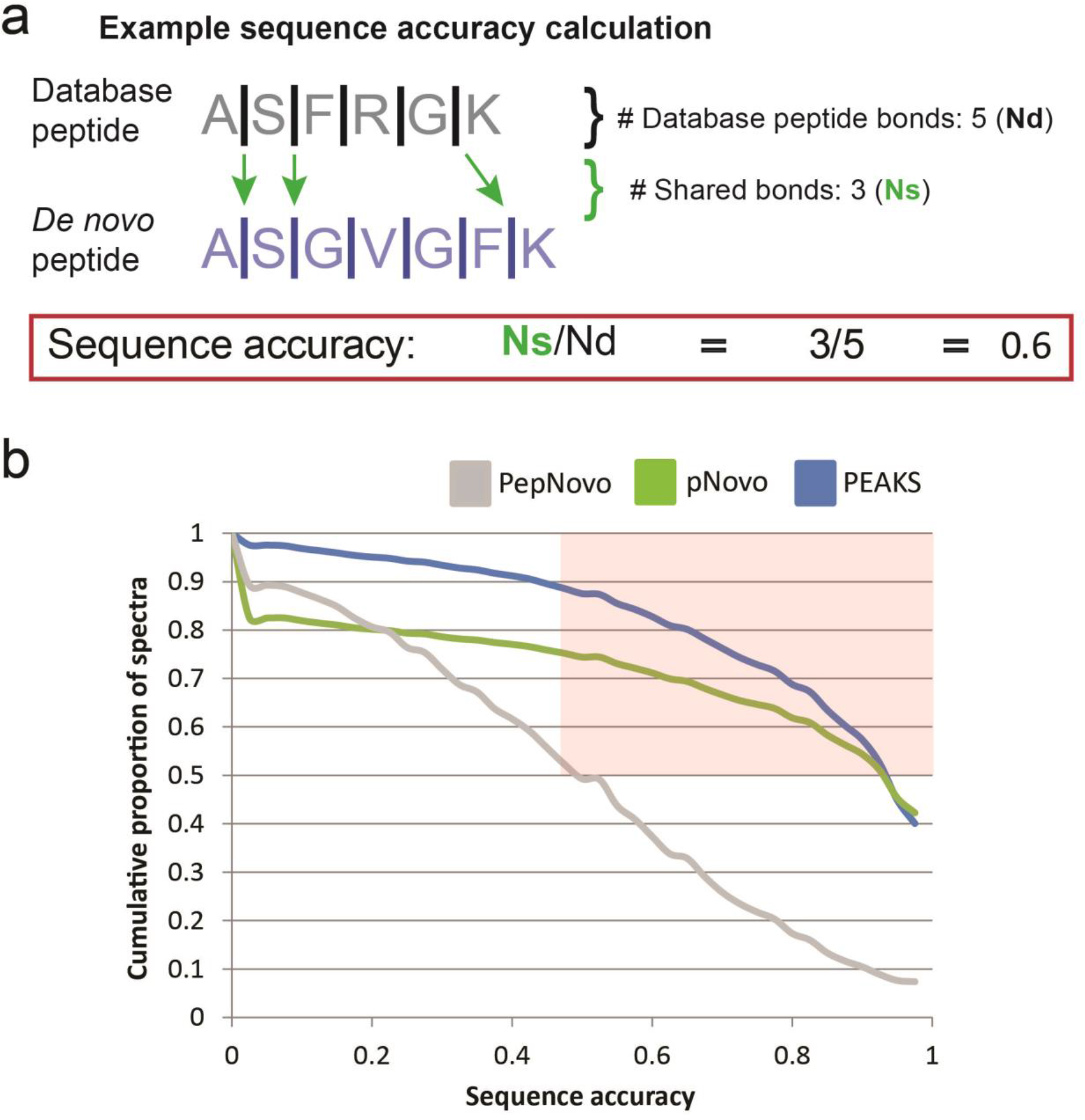
*De novo* sequencing algorithms yield mostly correct interpretations of most input spectra. **a)** Example calculation of sequence accuracy – the proportion of peptide bonds shared between a high-confidence peptide identification and the corresponding *de novo* peptide interpretation^9^. **b)** Cumulative proportion of spectra exceeding a given sequencing accuracy threshold (x-axis) for three *de novo* sequencers, PEAKS, PepNovo, and pNovo, as benchmarked on the A375 proteomic dataset (Fig. 1). PEAKS demonstrated the best performance overall.

**Supplementary Fig. 3.**
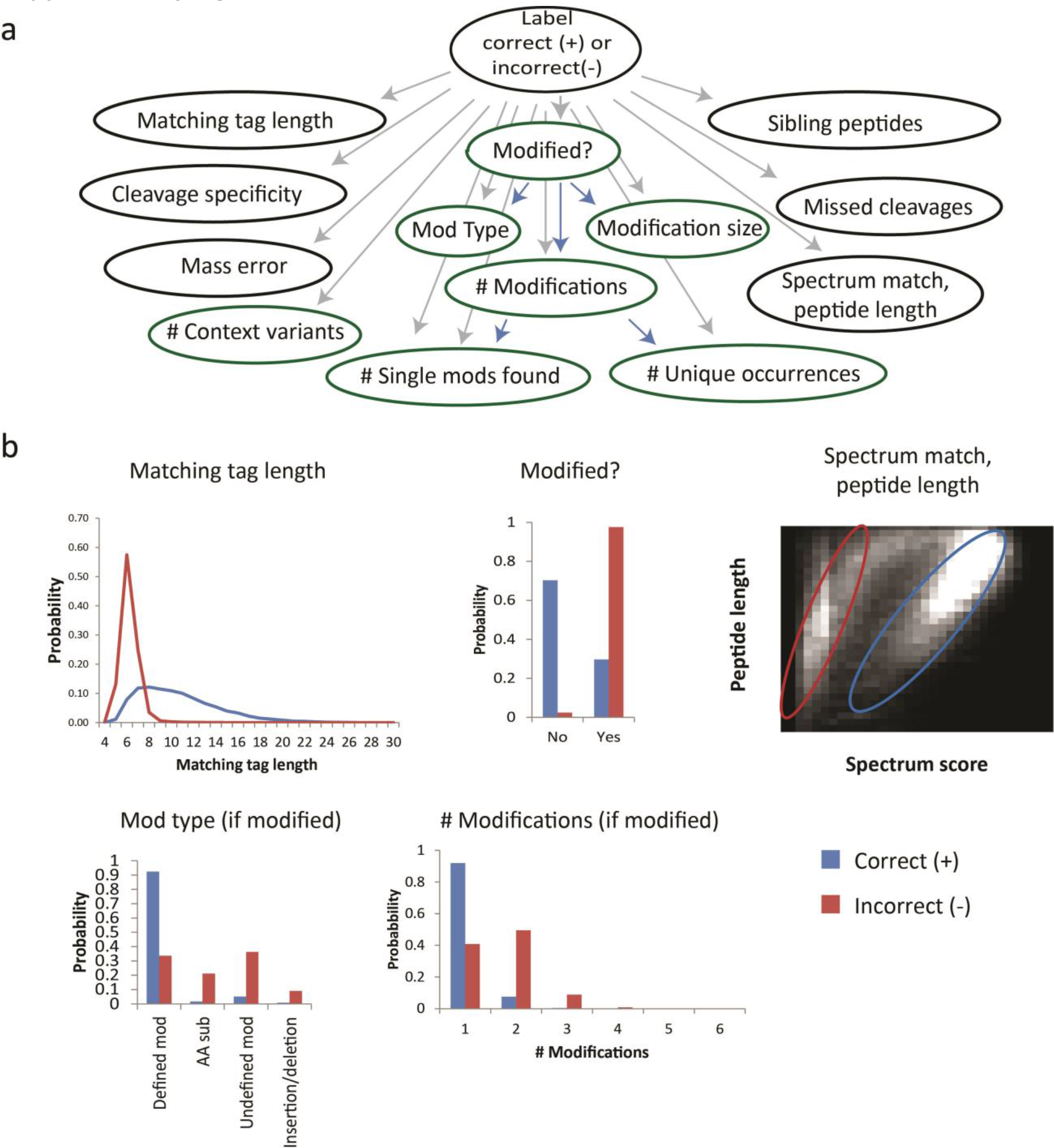
Hierarchical Bayes model description. **a)**Bayes model used for fitting correct (+) and incorrect (-) peptide-spectrum match distributions. Grey arrows indicate dependencies between model attributes and the distribution being trained. Blue arrows indicate dependencies between model attributes. Attributes in green oval specifically pertain to sequence modifications. Further details are provided in **Supplementary Note 3**. **b)** Example distributions for several model attributes derived from the A375 dataset (Fig. 1). Likelihood distributions were iteratively refined across multiple measurement dimensions using expectation-maximization (EM).

**Supplementary Fig. 4.**
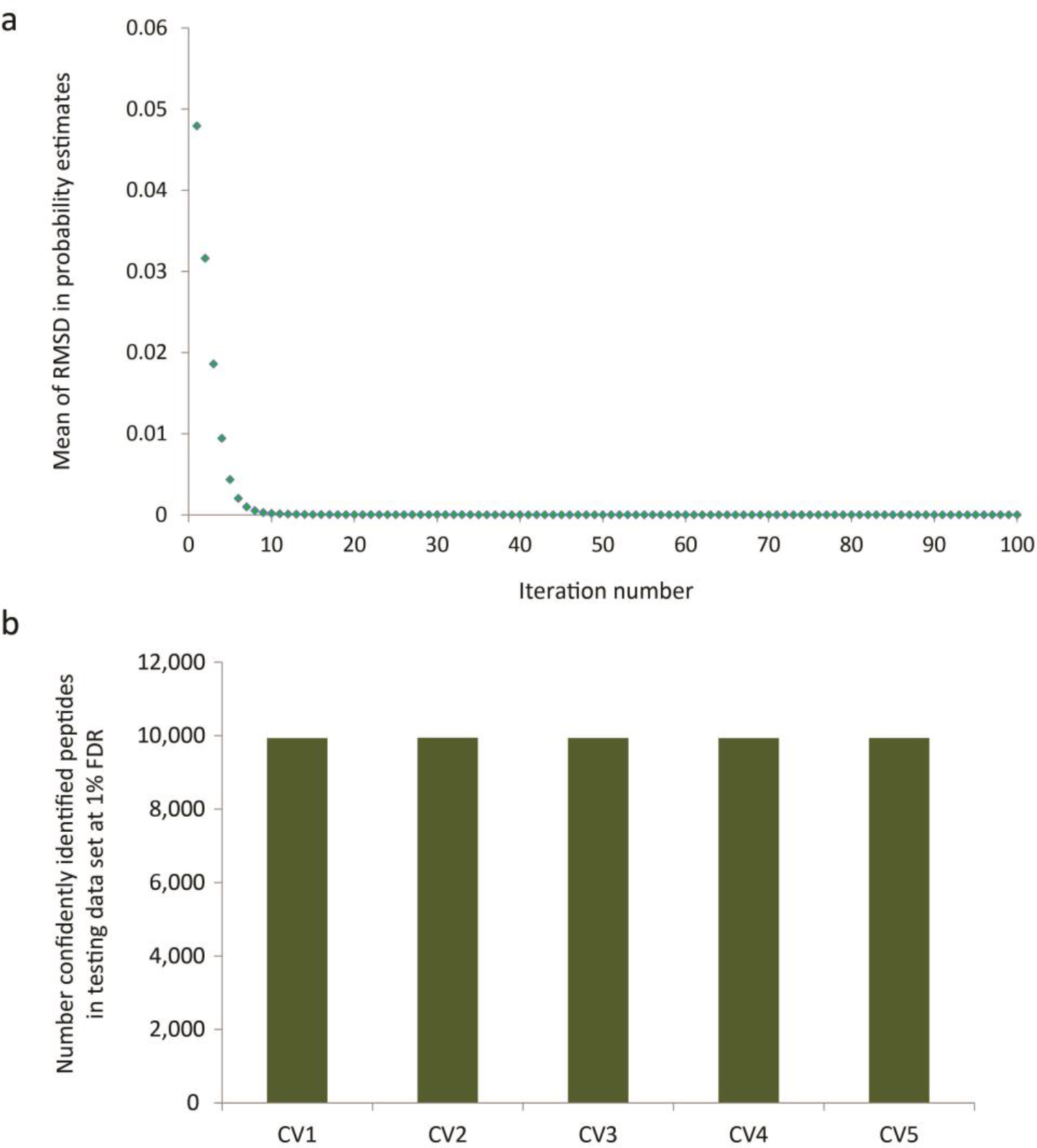
Expectation Maximization-estimated false discovery rate estimations are robust. **a)**Randomized starting model guesses for expectation-maximization-based training of the hierarchical Bayes model rapidly converged, and yielded highly consistent probability estimates. **b)** Five-fold cross-validation (CV) demonstrated that training the EM-optimized hierarchical Bayes model did not substantially affect the returned set of confidently identified spectra, when each model was tested on a dedicated test spectra set. Further details can be found in **Supplementary Note 3**.

**Supplementary Fig. 5.**
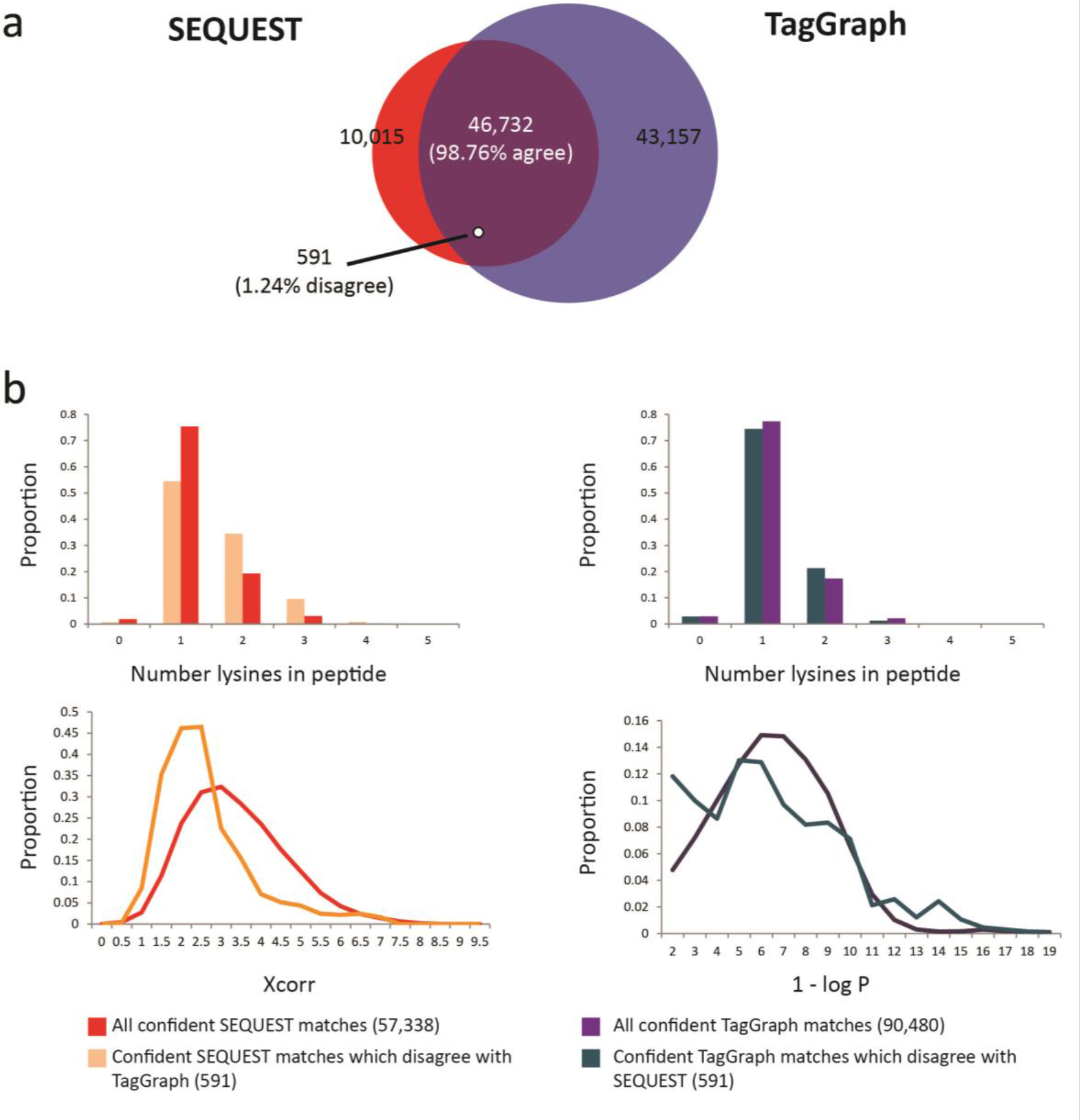
Conflicting high-confidence peptide-spectrum matches strongly favor TagGraph interpretations over SEQUEST. **a)**98.76% of 47,323 PSMs for which both TagGraph and SEQUEST return a high-confidence result (1% estimated FDR) agree, consistent with an estimated 1% FDR for both algorithms. PSMs were derived from the A375 dataset. **b)** Of the remaining 1.24% of PSMs for which SEQUEST and TagGraph disagree, TagGraph score and peptide missed cleavage distributions were more consistent with high-confidence identifications.

**Supplementary Fig. 6.**
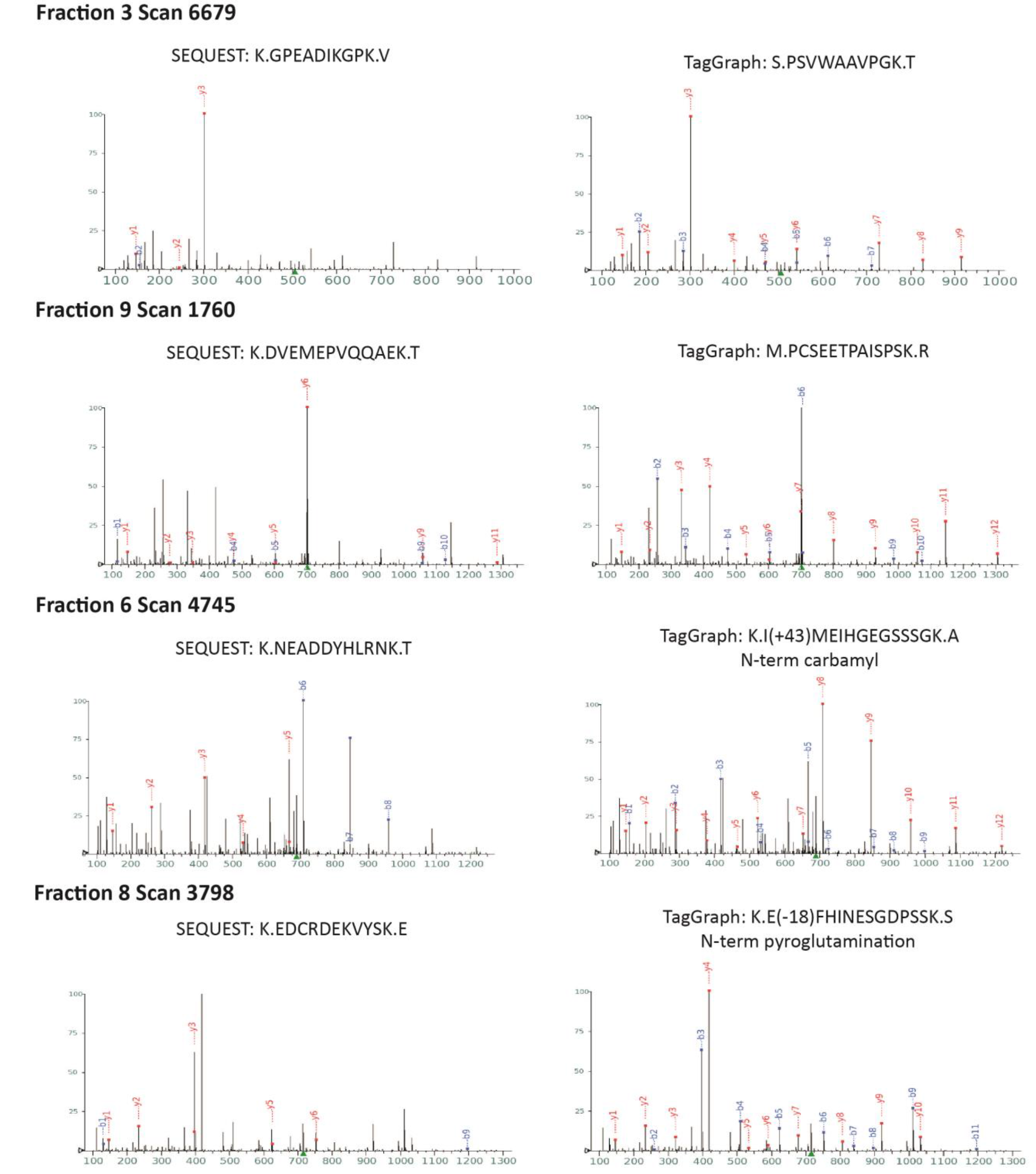
Examples of TagGraph-assigned peptide-spectrum matches that conflict with high-confidence SEQUEST assignments. Representative spectra demonstrating superior fragment ion assignments made by TagGraph for peptides more consistent with LysC digestion than the conflicting peptides SEQUEST assigned to the same spectra. Both results were assigned scores consistent with a 1% FDR on the A375 dataset with respect to each set of search results.

**Supplementary Fig. 7.** *[See file TG_Figure_S07_Search_Algorithm_TypeI_II_Errors.pdf]*

**Supplementary Fig. 7.** Examples spectra depicting “Case 1” (modification mislocalization) and “Case 2” (incorrect peptide sequence) interpretation errors, as defined Supplementary Note 2.

**Supplementary Fig. 8.**
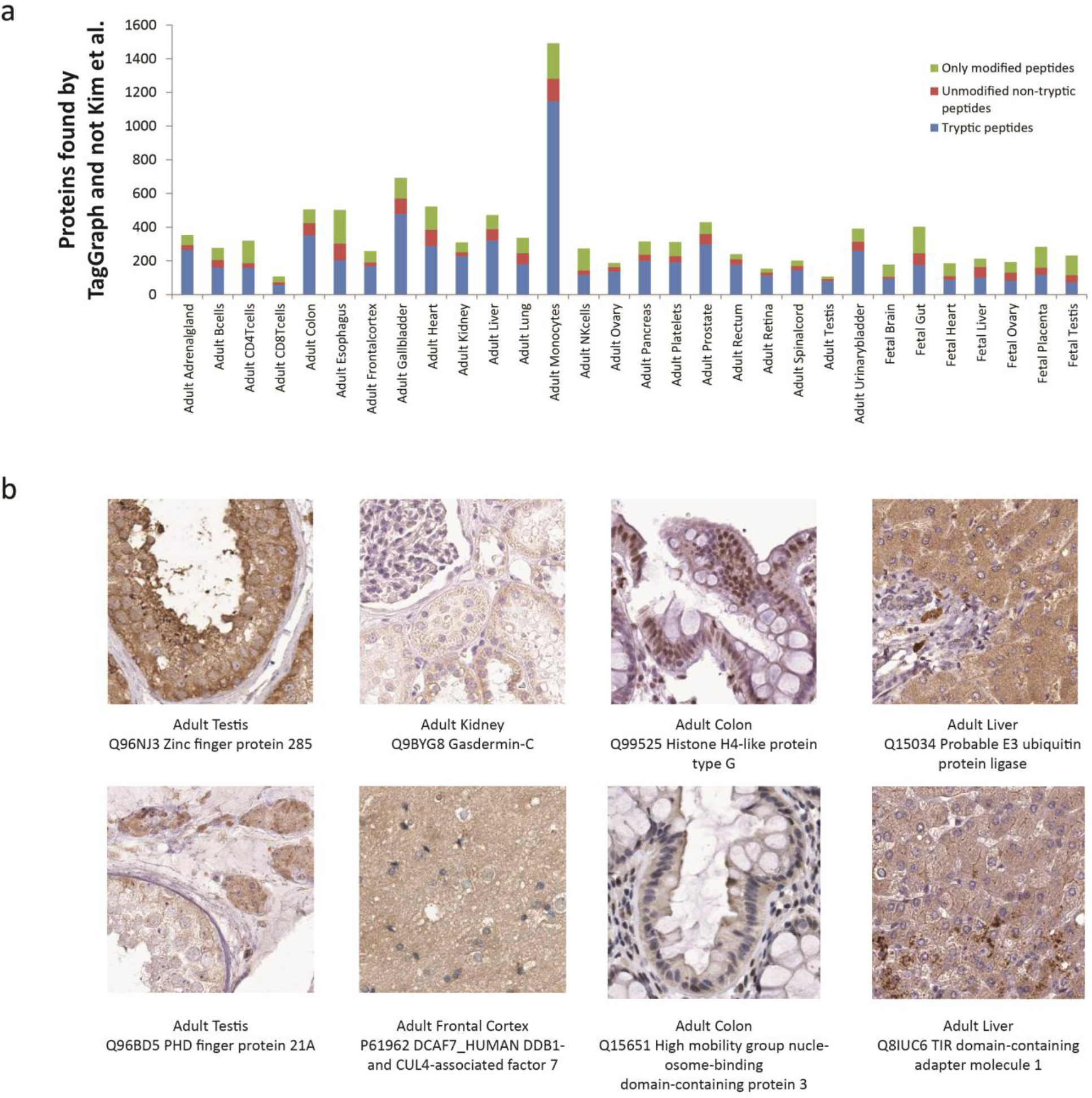
Increased proteome coverage by TagGraph relative to Kim *et al*. **a)**The number of proteins identified by TagGraph and not Kim et. al. are shown for each tissue examined in this dataset. Identified proteins were assigned one of three categories: (i) proteins with any unmodified tryptic peptides mapped to them, (ii) proteins with unmodified non-tryptic peptides mapped to them and no unmodified tryptic peptides mapped, and (iii) proteins with only modified peptides mapped to them. Proteins were designated as identified in the Kim et. al., analysis if at least one peptide was mapped to them, and proteins were designated as present in the TagGraph analysis if their normalized spectral abundance factor (NSAF) was greater than zero. We attribute the pronounced spike protein identifications from the Adult Monocytes tissue to a procedural error made by the study’s original authors: We found that the pepXML-formatted search result file corresponding with ‘bRP_Elite’ analysis, which we downloaded from the PRIDE database (PXD000561) was identical to the ‘bRP_Velos’ pepXML file. The raw data files corresponding with these two conditions were clearly distinct, and were used as input to TagGraph. This spike in identifications can only partially be attributed to TagGraph’s enhanced identification capabilities. **b)** Immunostaining images taken from ProteinAtlas^24^ for select proteins identified by TagGraph and not Kim et. al., validates TagGraph-specific protein predictions.

**Supplementary Fig. 9.** *[See file TG_Figure_S09_SyntheticPeptides.pdf]*

**Supplementary Fig. 9.** Synthetic peptides confirm novel post-translational modifications and other unexpected sequence variants TagGraph measured from draft map of the human proteome.

**Supplementary Fig. 10.**
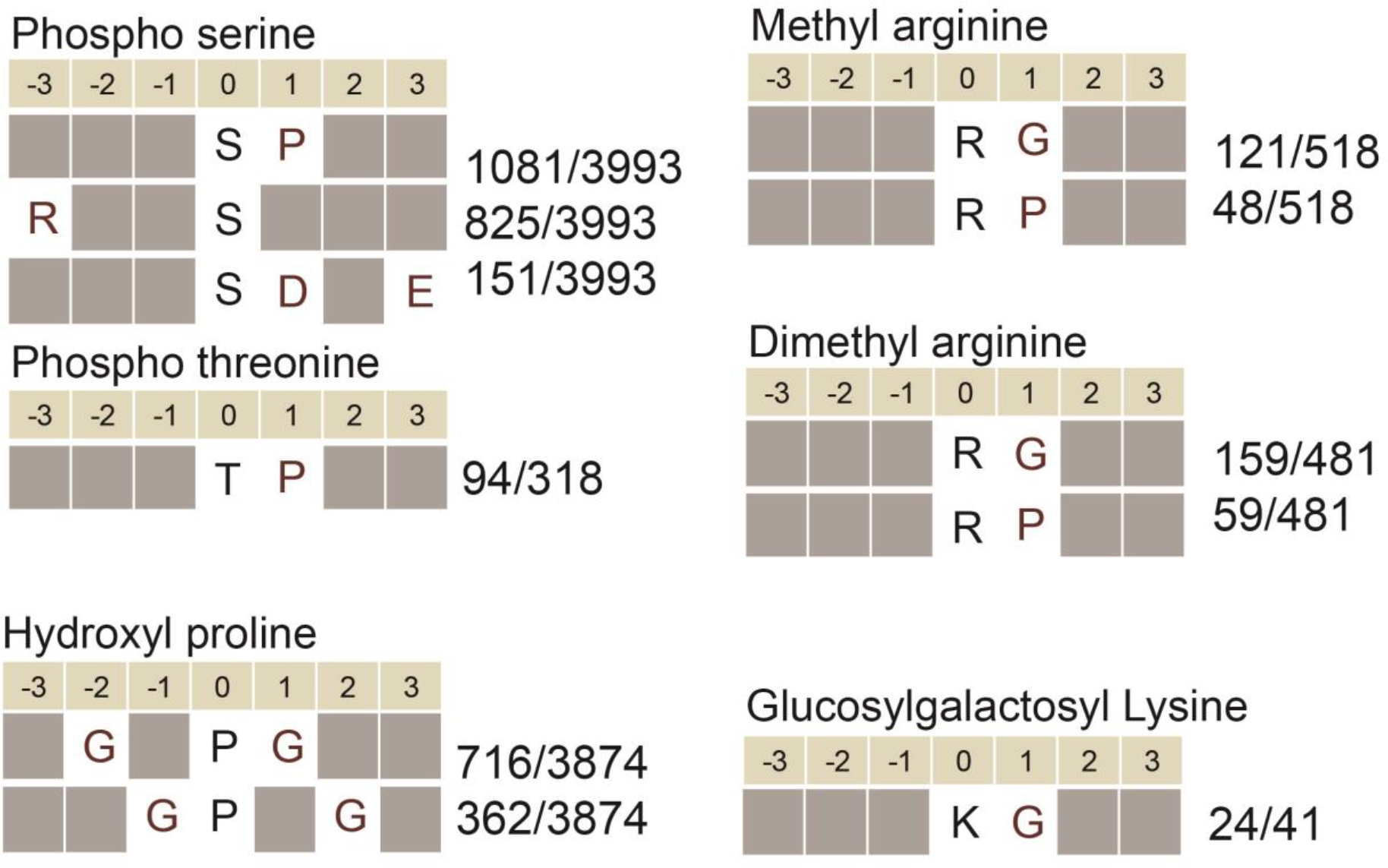
Motif-X analysis for known PTMs reveals known and novel substrate motifs. Motifs identified by the Motif-X algorithm^25^ surrounding several abundant PTMs. This analysis recovers known motifs for phosphoserine, phosphothreonine, dimethyl arginine, and methyl arginine PTMs, and predicts new motifs for the less well-characterized proline hydroxylation and lysine glucosylgalactosylation PTMs. Fraction indicates the number of times the indicated motif was identified out of the total number of modification sites entered into the Motif-X algorithm.

**Supplementary Fig. 11.**
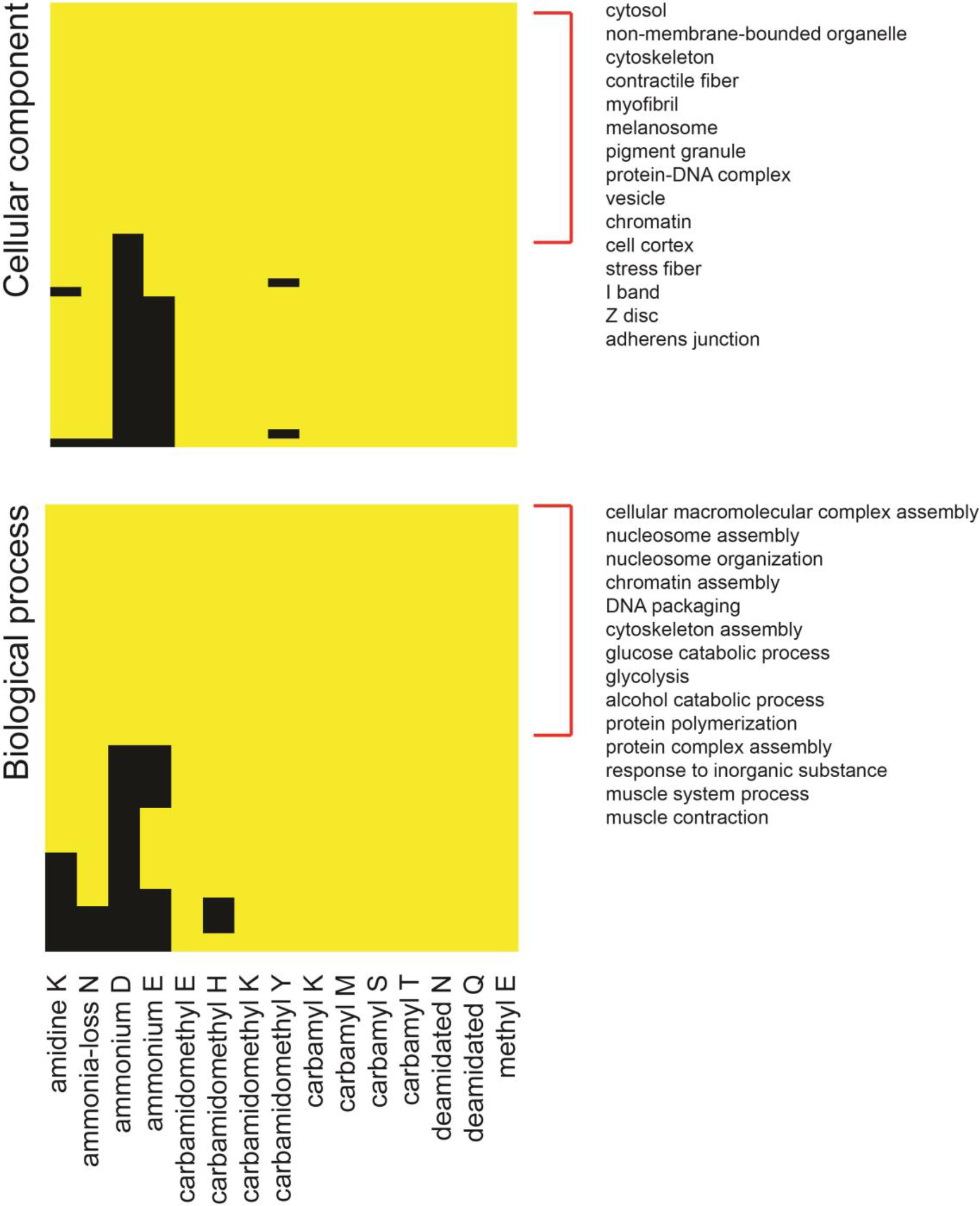
Accounting for ontologies enriched among post-isolation modifications. Gene ontology enrichment analyses of PTM-bearing proteins may be biased by mass spectrometers’ tendency towards identifying modified peptides from highly abundant proteins. Consequently, some ontologies could reach statistical significance based on protein abundance alone, rather than PTM-specific biological phenomena. To account for this, we identified significantly enriched ontologies (1% FDR, Benjamini-Hochberg corrected; yellow) among proteins bearing any of 15 abundant post-isolation modifications. Because these modifications should not have any inherent biological relevance, any ontology enriched among these post-isolation modifications were deemed false (red brackets), and removed from the analysis presented in Fig. 3b.

**Supplementary Fig. 12.**
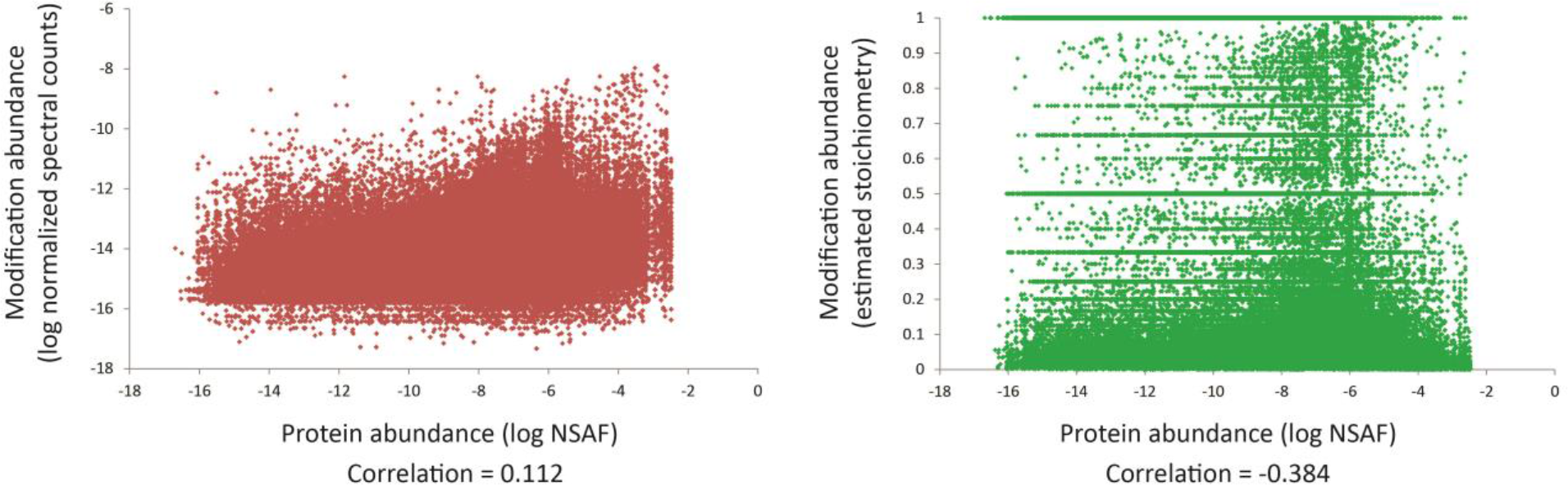
Modification abundances and stoichiometries are not correlated with protein abundances. Scatterplots of protein normalized spectral abundances factor (NSAF) with modification stoichiometry (left) or modification normalized spectral counts (NSC, right). In both cases, modification abundance did not correlate with protein abundance.

**Supplementary Fig. 13.**
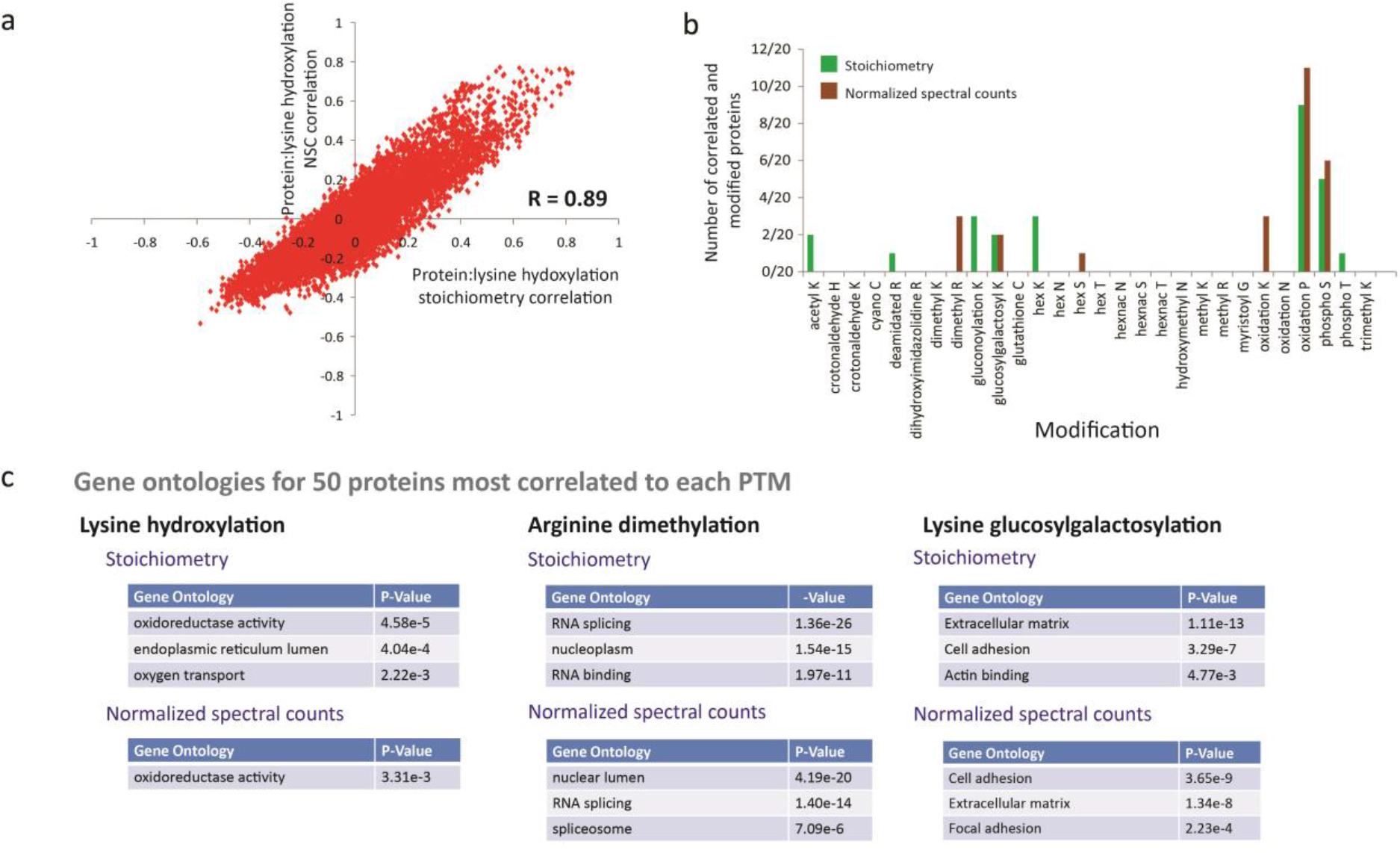
Proteins that correlate with PTM substrates share functional properties. **a)**Expression level (NSAF) profiles for 15,747 proteins spanning 30 tissues were correlated with averaged PTM profiles across the same tissues, using either stoichiometry or normalized spectral counts (NSC). The representative scatter plot shown here for lysine hydroxylation indicates the extent to which each protein’s tissue profile (points) correlates with lysine hydroxylation across the 30 tissues as measured by estimated stoichiometry (x-axis) or total abundance (y-axis). These data show that the two PTM quantification methods are broadly similar. However, protein correlation ranks may differ greatly between the two quantification methods. Thus, both can produce complementary but similar sets of highly correlated proteins.**b)** Protein-PTM correlations generally did not indicate specific modified substrates. A protein’s abundance could correlate with a particular PTM because it regulates or directly catalyzes the PTM’s formation on its substrate. Alternatively, proteins could be correlated with a modification because they are themselves heavily-modified substrates of the PTM. Kinases, which both catalyze phosphorylation events and are themselves highly phosphorylated, would be expected to be examples of both conditions, for example. By contrast, collagens would be examples of the latter condition, as abundant proteins in certain tissues that carry a highly degree of hydroxylated prolines. To evaluate these possibilities, we first identified the 20 proteins that most highly correlated with each of the 28 PTMs shown here, as computed using either modification NSC or stoichiometry. Of these, we plotted the number of proteins that were also modified by the indicated PTM. For the most part, however, PTMs were not identified on the same proteins to which they were most highly correlated, suggesting that they may be candidate regulators of PTM transfer. **c)** Enriched gene ontologies for the top fifty most correlated proteins for several PTMs suggests either enzymatic activity (i.e., oxidoreductase activity is known to be required for lysine hydroxylation to occur) or common functional activity (i.e., arginine dimethylation is known to be enriched in RNA splicing proteins, Fig. 3b). As demonstrated in part b, these proteins are themselves not substrates of the PTM of interest. Thus, these ontologies further suggest functional relationships between PTMs and proteins which are highly correlated with them.

## SUPPLEMENTARY TABLES

**Supplementary Table 1. TagGraph, PEAKS PTM, Byonic, ModA, and Open search results per mass spectrum from A375 cell line data set**

*[NB: this is a 92 Mb data file]*

a. Fraction number from high-pH reversed phase concatenated fractionation (see **Methods**)
b. Number of the MS2 fragmentation scan
c. Charge state of the precursor ion that gave rise to the indicated MS2 scan
d. Computed peptide’s mass, based on its amino acid sequence and any additional modifications
e. Inferred singly-charged ion mass, based on observed precursor ion’s m/z ratio and charge
f. Parts-per-million mass deviation between observed and theoretical peptide masses
g. Probability of indicated peptide identification being correct, as computed by search algorithm.
h. Log-transformed, inverse probability of indicated peptide identification being correct, as computed by the expectation maximization-optimized Bayesian network: -log10(1-p)
i. Peptide sequence from the input FASTA sequence database, noting the amino acids flanking the peptide, appearing outside the periods.
j. TagGraph-resolved peptide sequencing, noting deviations from the database sequence with a “-“
k. Modifications assigned to TagGraph-resolved peptide: Nested series are of the format: ((’*Mod1 name from Unimod if exists’*, *Mod1 delta mass from Unimod if it exists*, *Mod1 delta mass vs. Unimod if exists*), (*Mod1 target amino acid from Unimod if exists*, *Mod1 target amino acid location on peptide from Unimod if exists*), *indexed location of Mod1 on peptide sequence counting from zero*), ((*’Mod2 name from Unimod if exists’…*, *indexed location of Mod2 on peptide sequence counting from zero*),(…)]
l. List of proteins from FASTA sequence database containing indicated peptide
m. Score assigned to identified peptide by search algorithm
n. Peptide sequence, indicating position of modification within parentheses.
o. Inferred name and specificity of modification indicated in (n)
p. Example protein containing indicated peptide sequence
q. modified amino acid and rounded mass of corresponding modification
r. Peptide sequence, indicating position of modification with numerical modification representation immediately following modified residue
s. Deviation (Da) between observed and theoretical (unmodified) peptide masses

**Supplementary Table 2 High-confidence TagGraph results per mass spectrum from human proteome data set** *[NB: this is a 10.2 Gb data file]*

a. Tissue from which mass spectrum was derived
b. Acquisition method for mass spectrum, using the format [separation method (SDS-PAGE (“Gel”) or high-pH reversed phase (“bRP”))]_[mass spectrometer (LTQ Orbitrap Velos (“Velos”) or Orbitrap Elite (“Elite”)]
c. Fraction number from high-pH reversed phase concatenated fractionation (see **Methods**)
d. Number of the MS2 fragmentation scan
e. Retention time (minutes) of the indicated MS2 scan
f. Charge state of the precursor ion that gave rise to the indicated MS2 scan
g. Inferred singly-charged ion mass, based on observed precursor ion’s m/z ratio and charge
h. Computed peptide’s mass, based on its amino acid sequence and any additional modifications
i. Parts-per-million mass deviation between observed and theoretical peptide masses
j. Probability of indicated peptide identification being correct, as computed by the expectation maximization-optimized Bayesian network.
k. log-transformed, inverse probability of indicated peptide identification being correct, as computed by the expectation maximization-optimized Bayesian network: -log10(1-p)
l. peptide sequence from the input FASTA sequence database, noting the amino acids flanking the peptide, appearing outside the periods.
m. TagGraph-resolved peptide sequencing, noting deviations from the database sequence with a “-“
n. Modifications assigned to TagGraph-resolved peptide: Nested series are of the format: ((’*Mod1 name from Unimod if exists’*, *Mod1 delta mass from Unimod if it exists*, *Mod1 delta mass vs. Unimod if exists*), (*Mod1 target amino acid from Unimod if exists*, *Mod1 target amino acid location on peptide from Unimod if exists*), *indexed location of Mod1 on peptide sequence counting from zero*), ((*’Mod2 name from Unimod if exists’…*, *indexed location of Mod2 on peptide sequence counting from zero*),(…)]
o. list of proteins from FASTA sequence database containing indicated peptide
p. peptide sequence derived from *de novo* sequencing
q. score assigned to *de novo* sequenced peptide by *de novo* algorithm

**Supplementary Table 3** Normalized spectral abundance factor (NSAF) for all proteins TagGraph measured from Kim et al data set.

**Supplementary Table 4** All modifications found by TagGraph and their corresponding number of peptide-spectrum matches and unique peptides. Sites with at least 20 spectral counts were reported in modification counts reported in text.

**Supplementary Table 5** Normalized spectral count (NSC) for all modified sites TagGraph measured from Kim et al data set.

**Supplementary Table 6** Table S6: All enriched ontologies and corresponding p values for 22 noteworthy PTMs (biological process and cellular compartment)

**Supplementary Table 7** All mono- and dimethylation sites with normalized spectral counts (NSC) and corresponding protein abundances (NSAF).

**Supplementary Table 8** Estimated stoichiometry for all modified sites TagGraph measured from Kim et al data set.

**Supplementary Table 9** PTMs TagGraph assigned to five representative histone isoforms, aligned with prior histone PTM compilations.

**Supplementary Table 10** Supplementary Table 10. Significant overlap between COSMIC cancer mutations and TagGraph proline hydroxylations

## SUPPLEMENTARY NOTES

### Supplementary Note 1 TagGraph

**A. FM-index procedure.** The FM-Index^26^ implementation used in TagGraph is a fork of an existing open source implementation (https://github.com/mpetri/FM-Index). To create an FM-index of the human proteome ^12^, we first concatenated all sequences in the FASTA-formatted protein database into a flat sequence file. A separate database of protein start offsets is maintained for retrieval of protein annotations. A Burrows-Wheeler Transform^27^ was then applied to the flat sequence file for fast substring search, the results of which were compressed using an RRR Wavelet Tree ^28^ for efficient in-memory storage. The number of occurrences of a candidate sequence pattern in an index can be computed in O(N) time, where N is the length of the input pattern. The locations of the input pattern in an index can be retrieved in O(M) time, where M is the number of occurrences of the input pattern in the index.

**B. TagGraph algorithm.** The TagGraph algorithm takes as input a set of high-resolution MS/MS spectra, a corresponding set of *de novo* sequence interpretations, and an FM-Index constructed from a protein FASTA file (**Supplementary Note 1a**). The algorithm then generates and ranks a list of candidate peptide-spectrum matches for each input MS/MS spectrum with respect to the indexed protein sequences.

TagGraph first computes the maximum matching substring between an input *de novo* sequence and the FM-index, then retrieves all candidate protein sequences from the index which contain this substring. For each candidate, TagGraph first computes all amino acid dimers which match between the *de novo* sequence and protein sequence. Contiguous dimers are then merged into longer sequence substring “tags,” which are then input as nodes into a directed acyclic graph.

Edges are drawn between any two nodes to represent regions in which the *de novo* sequence and matching candidate peptide sequence disagree. This disagreement could be due to a *de novo* sequencing error or the presence of a sequence variant, post-isolation modification, post-translational modification, or previously uncharacterized mass shift, in the peptide which gave rise to the source spectrum. Each edge is annotated with its possible interpretations and weighted based on a heuristic scoring scheme designed to weight more likely explanations more highly (Supplementary Fig. 1). The top scoring peptide matches for each candidate are retrieved from the graph as the top scoring paths between a set of start and end nodes ^29^.

These start and end nodes represent potential start and end sites for a peptide interpretation in the candidate matching protein. The scored evidence supporting a peptide-spectrum interpretation from its start to its end nodes is referred to as its **path score**.

Once identified, each candidate peptide match is scored against the observed MS/MS spectrum using a probabilistic model to derive its **spectrum score**. This model scores all fragment ions that support the peptide identification according to the relative likelihood of measuring the observed fragment ion intensity versus random chance (Poisson) (Supplementary Fig. 1). The probabilistic fragmentation model described above was trained on a library of 20,000 high confidence in-house generated HCD MS/MS spectra, collected from peptides unrelated to the current study.

To improve TagGraph’s ability to discriminate true PTMs over confounding isobaric interpretations (e.g., two phosphorylation events on neighboring serines versus the less likely explanation of pyrophosphorylation on a single serine, Supplementary Fig. 1), the algorithm creates a second set of candidate peptide-spectrum matches to explore alternate, isobaric modifications learned from the input dataset: 1) All combinations of mass shift and corresponding amino acid are tallied from the initial set of candidate peptide spectrum matches, to create a mass shift – amino acid frequency matrix. 2) Each candidate peptide’s mass shift is evaluated with respect to the frequency matrix generated in (1). 3) If the peptide’s mass shift is equal to an existing modification corresponding with the same peptide assigned to a different MS/MS spectrum, is more prevalent in the entire dataset than the existing modification, and has a valid modifiable amino acid on the unmodified peptide sequence, then a peptide with this alternate modification is added as a match candidate to the corresponding spectrum. 4) If the mass of a candidate modification can be explained by the sum of two modifications represented in the list learned in (1), an additional peptide carrying these two PTMs will be added as a match candidate to the corresponding spectrum. This will occur, however, only if the expected number of peptides with this combination of modifications in the dataset is greater than one, and if both modifications have valid sites on the unmodified candidate peptide sequence. 5) All primary and secondary match candidates are assigned a path score and spectrum score as described above. 6) These are combined and ranked with the existing peptide-spectrum matches from the first round of candidate generation (Supplementary Fig. 1).

### Supplementary Note 2. Target-decoy error estimation is poorly suited to unrestricted search results

The target-decoy validation methodology is readily applicable to search results generated by conventional database search engines. Peptides identified by unrestricted search engines like TagGraph pose several challenges that were not anticipated in our initial description and of this error estimation tool, and which violate the major assumptions we proposed^30–32^. For target-decoy to accurately estimate false discovery rates, the set of decoys must be chosen such that incorrect identifications have an equal chance of matching either the target or decoy databases. For conventional database search, choosing a decoy database composed of the reversed counterparts of sequences in the target database largely satisfies this criterion. However, even relatively simple searches permitting just one variable modification (e.g., phosphorylation), secondary validation methods (e.g., Ascore ^33^ are needed to measure the modification’s site localization accuracy. This issue arises because the assumptions underpinning target-decoy are violated: the likelihood of a correct (target) peptide bearing an incorrectly-localized modification matching a given MS/MS spectrum is far greater than an incorrect, decoy peptide. As we will demonstrate below, this problem is greatly compounded when considering hundreds of modifications simultaneously (as Peaks PTM does), and becomes exponentially worse still when allowing arbitrary mass modifications and no protease specificity. In all cases, errors pertaining to the identity and localization of the annotated modifications and the location of the peptide sequence in the proteome cannot be accurately estimated using target-decoy with reversed sequence decoys.

#### Case 1: Improper modification annotation and localization

This case is the most common source of error in unrestricted database search, and is not addressed by target-decoy based validation. When considering a peptide sequence with a potential modification, an unrestricted search algorithm must score all possible localizations of that modification on the peptide sequence. The best, or highest scoring localization is often returned, although other localizations could be considered as well. Every residue in the peptide as well as the peptide’s N- and C-termini serve as potential modification sites.

Algorithms which consider large numbers of known residue-specific modifications but not undefined mass shifts, such as Peaks PTM and Byonic (not in wildcard mode) are still confronted by extremely large numbers of possible localizations. For instance, according to Unimod, methylation (+14.01565 Da) can be localized to the peptide N-terminus or C-terminus, as well as the amino acids C,H,K,N,Q,R,D,E,S, and T. Furthermore, many modifications in Unimod have similar, near-isobaric masses, increasing the set of potential localizations beyond those suggested by the modification itself.

Another potential source of FDR estimation error lies in determining the number and types of modifications assigned to a peptide. When considering a peptide with a known mass shift corresponding to a limited set of potential modifications, an algorithm must decide whether the mass shift corresponds to a single modification, two modifications in combination, or three or more. Even when considering only well-defined modifications, the combined mass of two modifications often equals the mass of third modification – two phosphorylations equal the mass of one pyrophosphorylation; two methylations equal the mass of one dimethylation, etc. There are also a considerable number of cases in which a single parent mass could correspond to various combinations of different modifications. For instance, the mass 86.00 Daltons could correspond to two carbamylations or one acetylation and one carboxylation. In the case of arbitrary mass modifications, a given mass shift could correspond with an effectively infinite range of modifications and modification combinations. The enormity of the set of possible modifications also limits the effectiveness of the open search method ^16^, necessitating user-supervised follow-up analysis to identity the modification corresponding with a measured mass shift, and to localize it on the returned peptide sequence.

Due to the abundance of candidates considered by the search engine, errors in modification annotation are common in all unrestricted search methods. These errors can arise from low-quality mass spectra, resulting from noise, incomplete fragmentation, co-isolated precursor ions, or other confounding features. They can also arise from the search engine’s configuration, such as internal scoring functions that inappropriately give extra weight modifications believed to be more likely *a priori,* or because the algorithm did not consider the true modification annotation as a candidate. In general, peptide-spectrum matches with incorrect modification annotations share many of the same b- and y-ions as the correct modification annotations. Thus, it is far more likely for a peptide to match an incorrect modification annotation than a reversed decoy sequence. A few examples of such errors produced by previously published unrestricted search algorithms are provided (**Supplementary Figure 7**).

Rare but biologically important PTMs can often occur at rates orders of magnitude lower than the most common post-isolation modification in a dataset. Without an accurate method to estimate FDRs of modification annotations, systematic algorithm-dependent errors are subject to being mistaken for these kinds of rare peptide-spectrum matches, limiting the utility of unrestricted modification searches for finding them. TagGraph’s hierarchical Bayesian model addresses this issue by making no assumptions about types of modifications present in the dataset *a priori* (or about any other attribute of the dataset). Thus, the validation model learns the distribution of modifications present, and weights its confidence in a particular modification annotation against other attributes of the peptide-spectrum match, such as the evidence provided in the spectrum, localization of the peptide sequence, and other attributes.

#### Case 2: Incorrect base peptide sequence

The large set of possible modifications an unrestricted search algorithm must consider also makes it possible for such algorithms to choose an incorrect base peptide sequence. This incorrect peptide sequence could originate from a protein other than the correct sequence (i.e., a sequence from a homologous protein containing a single amino acid polymorphism), or could be a slight deviation from the correct peptide sequence (i.e., with several amino acids added or truncated from the N- or C- terminus). In all cases, these incorrect base peptide sequence predictions can be reconciled with the MS/MS spectrum’s precursor mass through erroneous modification annotations. As above, these incorrect base peptide matches are more likely than decoy sequence matches to a spectrum since they often share many b- and y-ions with the correct peptide-spectrum match. A few examples of such errors are provided (**Supplementary Figure 7**).

The rate at which these types of errors occur is determined by the degree of sequence self-similarity present in the sequence database and the number of modifications considered. As the number of modifications considered increases, more base peptide sequences can be considered as candidate matches for a peptide spectrum match. Similarly, as sequence self-similarity increases (i.e., due to homology), more sequences similar to the true peptide sequence can be considered as candidates: mass shifts assigned to one or more modifications can make them consistent with the spectrum’s precursor mass, even with very high degrees of measurement mass accuracy.

#### Exacerbation of Case I and Case II errors when allowing arbitrary protease specificity

We have shown with specific examples and with global analysis that previously described unrestricted search algorithms are prone to the above errors cases. Furthermore, the rates of these errors greatly exceed what would be expected using target-decoy FDR estimation. This error underestimation problem could be worse for TagGraph, since it is the first to perform truly unconstrained search, with respect to protease specificity, and arbitrary types and numbers of modifications, and sequence variants. This larger search space raises the likelihood of matching an incorrect peptide sequence to an input spectrum by chance. We explore several of these possibilities below.

Peptides that are inconsistent with protease specificity can result from either in-source fragmentation or endogenous protease activity. Though often excluded from reported datasets, TagGraph was designed to readily detect them in either modified and unmodified forms (**Supplementary Note 1**). Spectra corresponding to such peptides can return high-scoring incorrect identifications with a protease-constrained unrestricted search (i.e., returning the nearest protease-specific peptide with a modification annotation --Open search is prone to this error). As such results must then be manually separated from true, modification-bearing peptides, an algorithm such as TagGraph which can correctly annotate these peptide-spectrum matches from the outset is highly desirable.

However, opening up the search space in this way poses substantial challenges in validation in addition to those presented above. Namely, the set of candidates which must be considered by the algorithm increases exponentially. For instance, consider an MS/MS spectrum for which the correct identification is the following peptide-spectrum match:

> …KAARE<u>E(+43)TDFCEEDSIEK</u>SS…

Where the underlined sequence indicates the returned peptide-spectrum match, placed in the context of its surrounding protein sequence in the database. The +43 indicates an N-terminal carbamylation. An algorithm which allows arbitrary mass modifications and no protease specificity must consider an effectively infinite number of candidates. However, unlike the case in which protease specificity is constrained *a priori*, the algorithm must also consider multiple base peptide sequences. For example, here are several incorrect annotations which could be candidate matches for the spectrum yielding the true match above:

> …KAAR<u>EE(-86)TDFCEEDSIEK</u>SS…
>
> …KAAR<u>E(-86)ETDFCEEDSIEK</u>SS…
>
> …KAAREE<u>T(+172)DFCEEDSIEK</u>SS…
>
> …KAAREE<u>T(+43)DFCE(+129)EDSIEK</u>SS…
>
> …KAARE<u>E(+43)TDFCEEDSIE(+128)</u>KSS…
>
> …KAAR<u>E(-86)ETDFCEEDSIE(+128)</u>KSS…

These annotations have similar predicted fragmentations to the correct peptide-spectrum match. As described in Case I and II above, this similarity makes them far more likely than a match to a decoy sequence in the case of a false match, rendering target-decoy unsuitable. Algorithms which constrain the set of modifications and protease specificity face the possibility of Case I and II errors on a subset of spectra, and commit them involuntarily by excluding the correct peptide-spectrum match from the set of candidates on another subset of spectra. By allowing arbitrary mass modifications and no protease specificity, TagGraph must be able to discriminate correct peptide spectrum matches from Case I and Case II errors for *every* spectrum.

We address this problem by using an expert-designed hierarchical Bayesian model (**Supplementary Note 3**) which can learn the specific attributes of correct and incorrect peptide-spectrum matches from each dataset individually, including the attributes of correct and incorrect modification annotations. This avoids problems present in the scoring functions of some previously published unrestricted algorithms, which make *a priori* assumptions on the likelihood of observing certain modifications in the dataset that are often untrue. Unlike target-decoy, this model is able to assign higher or lower confidence to a peptide-spectrum match based on its modification annotation, and consider this attribute in the context of the peptide localization, evidence in the spectrum, etc. The model is both flexible and extensible, enabling further refinement based on discriminatory criteria discovered to be useful in the future.

### Supplementary Note 3. Estimating peptide identification error through a hierarchical Bayes probabilistic model, optimized by Expectation Maximization

#### A. Overview

We developed a peptide identification error model that overcomes the limitations conventional target-decoy searching has for assessing modification-bearing peptide identifications. Our hierarchical Bayes model^7^ creates hypothetical structures for the probability distributions corresponding with observing a set of data features D given that the peptide interpretation is correct P(D|+) or incorrect P(D|-). D is defined as the set of peptide and fragmentation spectrum attributes that are represented by the hierarchical Bayes model (Supplementary Fig. 3a). These attributes were empirically chosen based on their utility in discriminating correct peptide-spectrum matches from incorrect ones. Considering both of the above probability distributions, we calculate P(+|D), the probability that any given peptide-spectrum interpretation is correct, using Bayes Theorem.

#### B. Model attributes

The following describe the general and modification-specific attributes used in our hierarchical Bayes model, as represented in Supplementary Fig. 3a.

##### General peptide attributes

*Spectrum Score*: The spectrum score of a candidate peptide-spectrum match, as defined in (*iii*), above.

*Peptide Length:* The length of an entire candidate peptide-spectrum match. This attribute is modeled jointly with the spectrum score due to the observation that the spectrum score is negatively correlated with peptide length (Supplementary Fig. 3b).

*Cleavage Specificity:* If a protease was used to digest the sample, this attribute encodes whether a peptide-spectrum match has full protease specificity, a nonspecific n-terminus, a nonspecific c-terminus, or both nonspecific termini. This attribute is ignored if no protease was used.

*Missed Cleavages:* If a protease was used to digest the sample, this attribute records how many missed protease cleavage sites are present in a peptide-spectrum match.

*Matching Tag Length:* The length of the maximal matching substring between the *de novo* peptide interpretation and the candidate peptide-spectrum match.

*Sibling Peptides*: Number of other unique peptides found from the same protein that produced the current peptide-spectrum match.

*Mass Error*: The difference in Daltons between the mass of a peptide-spectrum match and the predicted mass based on the peptide sequence and the set of its annotated modifications.

##### Modification-derived Attributes

*Modified?:* Categorical attribute denoting whether or not the current peptide-spectrum match has modifications with respect to the underlying database sequence.

*Maximum Modification Mass:* The mass (Da) of the largest modification assigned to a candidate peptide-spectrum match.

*Number Unique Occurrences:* The number of unique peptides observed in the dataset with the same set of modifications as the candidate peptide-spectrum match.

*Modification Type:* Categorical labeling of annotated modification as either an amino acid substitution, defined modification (i.e., present in Unimod but not an amino acid substitution), insertion/deletion, or undefined mass shift.

*Number of Modifications:* The number of modifications annotated on the peptide-spectrum match.

*Number of Context Variants:* The number of unique peptides present in the dataset with the same base peptide sequence as the candidate peptide-spectrum match. A unique peptide is defined by its primary amino acid sequence, all modifications it may contain, and the corresponding modification positions along the sequence.

*Number of Single Modifications Found On Same Context:* If a given peptide-spectrum match contains multiple modifications, this attribute records the number of distinct singly-modified forms of the same peptide found in the entire dataset.

#### C. Learning the distributions P(D|+) and P(D|-)

As the set of correct and incorrect peptide-spectrum matches is not known *a priori*, we use an expectation maximization algorithm to learn the distributions P(D|+) and P(D|-) from observed data features D. Learning these distributions is accomplished through two steps: a re-ranking step and a convergence step.

Several model attributes in the hierarchical Bayes model rely on quantities calculated from the set of all peptide-spectrum matches in a dataset. These attributes can significantly affect the probabilities P(+|D) of candidate peptide-spectrum matches for each spectrum. Thus, the estimate of P(+|D) within a given iteration of the expectation-maximization process can change both the attributes D for all peptide-spectrum matches and the optimal match for a particular spectrum relative to the previous iteration. We expect the rankings produced by the probabilities P(+|D) to be superior to the initial rankings TagGraph produces using the sum of the spectrum and path scores alone. Given that the estimate of P(+|D) changes with each iteration, we further expect more sensitive discrimination between correct and incorrect identification by allowing the rankings of candidates for each spectrum to shift as the estimate of P(+|D) increases in accuracy. This intuition informs the basis of the re-ranking step of model training. In this step, the estimates of P(D|+) and P(D|-) are calculated using the top scoring candidates for each spectrum per expectation-maximization iteration. The candidates for each spectrum are then re-ranked according to their corresponding probabilities P(+|D), and the estimates of P(D|+) and P(D|-) in the next expectation-maximization iteration are calculated based on this new set of top-ranked candidates. After 20 rounds of re-ranking, the top ranked candidates are fixed in position and the algorithm is allowed to converge on stable estimates of P(D|+) and P(D|-) during the convergence step. Model variance is calculated as the Euclidean distance between the vectors P(+|D) for all spectra between the current and previous iteration. Convergence of the model to near zero variance typically occurs within 100 iterations

Within each iteration, the distributions P(D|-) and P(D|+) are learned as follows: the spectrum score and peptide length attributes are fitted as a multivariate Gaussian using a maximum likelihood estimator, weighted by the probability estimates derived from the previous expectation maximization iteration. The remaining model attributes are discretized into bins: the probability of observing each bin B for a given attribute A for the (+) distribution in the current expectation-maximization iteration is calculated using the estimates of P(+|D) from the previous iteration according to the familiar formula^34^:

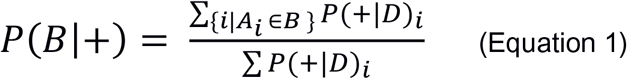

The formula for calculating attribute probabilities for the (-) distribution can be analogously generated using estimates of P(-|D). Before the first iteration, the EM algorithm is supplied with an initial guess for the parameters of the multivariate Gaussian describing the peptide length and spectrum score and for the distribution of the matching tag length attribute (Supplementary Fig. 3). Initial guesses for the parameters of both the correct (+) and incorrect (-) distributions are supplied. These guesses are used to populate initial estimates for the probabilities P(+|D) for each peptide-spectrum match in the dataset. These probabilities are then iteratively improved using the expectation maximization algorithm as described above. The spectrum-level probabilities P(-|D) can be readily converted to a global false discovery rate for a given set of spectra S using the following formula:

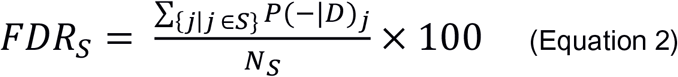

Where NS is the number of spectra in S. The large number of free parameters used to generate this model could be susceptible to overtraining with datasets of small size. However, we found that the algorithm converges onto accurate probability estimates for datasets of the size typically produced in modern proteomics experiments (Supplementary Fig. 4, **Supplementary Note 3D**).

### D. Evaluating EM model stability

We probed the robustness of the expectation-maximization based learning approach in two different ways, both using the cell line dataset described in Figure 1. First, the initial guesses used to seed the model training were randomly varied. The EM algorithm was run as described in section (iv), and the estimates of P(+|D) of all peptide-spectrum matches in the dataset were recorded for each iteration following the random initial guess. The root mean square deviation (RMSD) of each probability estimate was computed and averaged over the entire probability vector to derive the Mean RMSD over all initial guesses. This deviation asymptotically approaches zero with increased iterations, demonstrating that the final probability estimates are independent of the initial guess used (Supplementary Fig. 4a).

Second, to assess whether or not the EM model was susceptible to over-fitting, we employed five-fold cross validation. First, the dataset was randomly split into testing and training subsets, consisting of 10% and 90% of the peptide-spectrum matches, respectively. Slices consisting of 80% of the training dataset were randomly chosen and used to train the parameters of the P(D|+) and P(D|-) distributions. Estimates of the probabilities P(+|D) for the peptide-spectrum matches in the testing dataset were computed at each iteration using the models trained from each slice. The final probability estimates derived from the testing set were found to be independent of the slice used for training, demonstrating that the model learned general features of the data and did not over-fit to specific attributes of the randomly chosen subsets (Supplementary Fig. 4b).

